# Metal stress uncouples early pre-rRNA processing from ISR activation and reveals flexible checkpoints in human ribosome biogenesis

**DOI:** 10.1101/2025.07.14.664728

**Authors:** Anne Aldrich, Rachael Thomas, Michael Blower, Shawn M. Lyons

**Affiliations:** Department of Biochemistry and Cell Biology, Boston University Chobanian and Avedisian School of Medicine, Boston, MA 02118; Program in Genetics and Genomics, Boston University Chobanian and Avedisian School of Medicine, Boston, MA 02118; The Genome Science Institute, Boston University Chobanian and Avedisian School of Medicine, Boston, MA 02118

## Abstract

Ribosome synthesis is one of the most energy-intensive processes in a growing cell, consuming more than 60% of cellular energy reserves. As such, ribosome biogenesis is highly sensitive to stress to prevent costly expenditures under adverse conditions. Moreover, successful assembly requires precise stoichiometric balance between ribosomal proteins and ribosomal RNAs. Here, we define novel regulatory mechanisms of ribosome biogenesis under stress that reveal previously unrecognized aspects of rRNA maturation. We demonstrate that early pre-rRNA processing is particularly sensitive to stress induced by environmentally relevant heavy metals. Surprisingly, our analysis shows that 5′ and 3′ end processing can be uncoupled in human cells, with 3′ end cleavage occurring independently of 5′ end processing. We further show that classical inducers of endoplasmic reticulum stress suppress ribosomal protein synthesis without inhibiting rRNA transcription, leading to an imbalance between these essential components of ribosome assembly. This imbalance may exacerbate cellular stress and compromise proteostasis. Together, our findings uncover stress-specific checkpoints in ribosome biogenesis that link environmental exposures to disrupted nucleolar function and highlight new layers of regulation in human rRNA maturation.

## INTRODUCTION

*De novo* synthesis of ribosomes is one of the most energy-intensive processes in a growing cell, consuming over 60% of the cell’s energetic resources (1). To avoid wasteful expenditure, ribosome production is tightly regulated by growth and stress signaling pathways, ensuring that this process occurs only under optimal conditions (2). Each human ribosome is larger than 4 MDa and consists of four distinct non-coding ribosomal RNAs (28S, 18S, 5.8S, and 5S rRNAs) and 80 ribosomal proteins. Its assembly demands coordinated control over multiple processes: transcription of ribosomal protein mRNAs, their translation, synthesis of rRNA precursors, and the precise processing and chemical modification of those pre-rRNAs. Ribosome biogenesis engages all three nuclear RNA polymerases: RNA polymerase I and III transcribe rRNA precursors, while RNA polymerase II transcribes mRNAs encoding ribosomal proteins and most small nucleolar RNAs (snoRNAs), which guide rRNA modifications and processing (3,4). Most assembly steps occur within the nucleolus, a membraneless, liquid–liquid phase-separated (LLPS) organelle (5). The nucleolar structure is maintained by a high concentration of nascent pre-rRNA, which serves as both scaffold and substrate for processing into mature rRNAs. In parallel, ribosomal protein mRNAs are translated in the cytoplasm, and the resulting proteins are imported into the nucleus and assembled into pre-ribosomal particles within the nucleolus.

Maintaining a stoichiometric balance between ribosomal proteins and rRNA is essential for cellular homeostasis. Cells have evolved quality control mechanisms to rapidly degrade excess ribosomal proteins to prevent their toxic accumulation (6-9). In contrast, how cells adjust rRNA production in response to changes in ribosomal protein availability is less well understood. Depletion of individual proteins can lead to distinct pre-rRNA processing defects (10,11), a phenomenon linked to a class of diseases known as ribosomopathies, including Diamond-Blackfan Anemia (12,13). However, whether cells possess global mechanisms that monitor ribosomal protein levels and coordinate them with rRNA synthesis remains an open question.

The 18S, 28S and 5.8S rRNAs are initially transcribed as a long tricistronic precursor known as the 47S pre-rRNA (Figure 1). This precursor undergoes a series of endonucleolytic cleavages and exonucleolytic trimmings at defined sites to generate mature rRNAs incorporated into ribosomes. While most of the enzymes mediating these steps have been identified, often through initial discovery in yeast followed by characterization of mammalian homologs, the enzymes responsible for the earliest cleavages at the extreme 5’ and 3’ ends (sites A’/01 and 02) remain unknown. This gap reflects both a lack of sequence conservation between yeast and metazoans and limited genetic tools available to interrogate these initial processing events in human cells. Moreover, the regulatory coordination of A’/01 and 02 cleavage is poorly understood. In human cells, current models suggest that these two events occur simultaneously, rapidly converting 47S to 45S pre-rRNA, whereas in murine cells, 5′ end processing at A’/01 has been reported to precede 3′ end cleavage, producing a short-lived intermediate termed the 46S pre-rRNA. Whether such flexibility exists in human rRNA maturation and how it may be influenced by stress signaling, remains unresolved.

**Figure 1.**
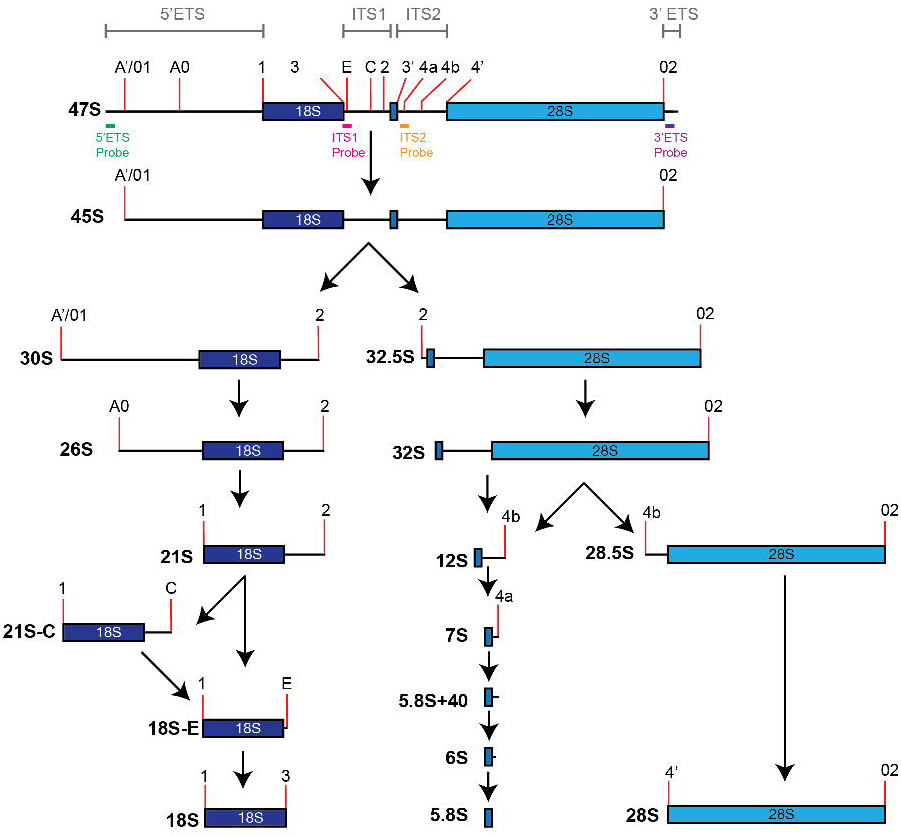
Simplified processing diagram of 47S pre-ribosomal RNA (pre-rRNA) transcript to mature 18S, 5.8S, and 28S rRNA. Location of Northern probes mapped to the 5’ externally transcribed spacer (ETS), internally transcribed spacer (ITS) 1, ITS2, and 3’ETS regions are noted on the 47S diagram. Major endonucleolytic and exonucleolytic processing sites are indicated.

Given its complexity and high metabolic cost, ribosome biogenesis is acutely sensitive to stress. rRNA transcription and translation of ribosomal protein mRNAs are both strongly regulated by mTOR signaling. Under nutrient-limiting or specific stress conditions, mTOR is inactivated, leading to coordinated repression of ribosomal protein translation via LARP1 or 4EBP (14,15) and downregulation of rRNA transcription through modulation or Pol I transcription factor TIF-IA (16). This dual regulation allows mTOR to synchronously inhibit ribosomal RNA and protein synthesis.

However, other stress signaling pathways do not appear to directly regulate both arms of ribosome production as directly. The integrated stress response (ISR) represents a second major mechanism that regulates protein synthesis during stress (17). ISR activation begins with one of four eIF2α kinases, HRI, PERK, PKR, GCN2, which phosphorylate the α subunit of eIF2, leading to global inhibition of translation including that of ribosomal proteins (18-21).

Our previous work demonstrated that some ISR-activating stresses also induce a defect in pre-rRNA processing at the A′/01 site, located near the 5’ end of the 47S precursor (2). We refer to this defect as "A′/01 stalling." However, the mechanistic relationship between ISR activation and A′/01 stalling is poorly understood. Our previous data suggest that while both pathways are triggered under stress, they may represent parallel but distinct branches of coordinated response that ensures rRNA synthesis is curtailed when ribosomal protein synthesis is impaired.

In this study, we define the molecular mechanisms underlying A′/01 stalling, reveal its independence from ISR activation and uncover metal specific mechanisms that reshape nucleolar function. Notably, we identify a novel 47S-C pre-rRNA species that accumulates when 3’ end cleavage occurs in the absence of 5’ end processing. This intermediate reveals, for the first time, that the earliest seps in pre-rRNA maturation can be uncoupled. We analyzed classical activators of the ISR, focusing on endoplasmic reticulum (ER) stressors that signal through PERK. Although these stressors robustly phosphorylated eIF2α and suppressed global protein synthesis, they failed to induce A′/01 stalling, indicating that this pre-rRNA processing defect is not universally correlated with ISR activation. We then turned our attention to a panel of environmentally relevant toxicants, sodium arsenite (NaAsO_2_), cadmium chloride (CdCl_2_) and chromium trioxide (CrO_3_), to determine if metal induced stress selectively activates this processing defect. All three compounds disrupted ribosome biogenesis, but through mechanistically distinct pathways.

## RESULTS

### AsO_2_ decouples 5’ and 3’ end processing resulting in accumulation of a novel pre-rRNA intermediate

AsO₂ is one of the most commonly used activators of the ISR, acting through activation of the heme-regulated inhibitor (HRI) kinase to phosphorylate eIF2α (18,22). Upon solubilization, AsO_2_ generates trivalent arsenic [As(III)], a highly reactive species and a major environmental toxin (23). To better understand how this stressor affects ribosome biogenesis, we analyzed pre-rRNA processing in cells exposed to AsO₂. We previously proposed that arsenite disrupts processing at the A′/01 site, leading to the accumulation of unprocessed 47S pre-rRNA (2), which retains both its 5’ and 3’ ends (Figure 1). While the enzymes responsible for A’/01 and 02 site cleavage remain unidentified (24), current models suggest that 5’ and 3’ end processing occur simultaneously in human cells, resulting in rapid conversion of 47S to 45S pre-rRNA. In contrast, studies in mice suggest that these events can occur sequentially, with 5’ end processing preceding 3’ end cleavage and giving rise to a short-lived 46S intermediate (25-27).

To gain better understanding of this process in humans and to determine how AsO_2_-induced A’/01 stalling affects 02 processing, we treated cells with AsO_2_ for 2 hours and pre-rRNA was analyzed by northern blotting using specific probes that detect 5’ ETS (Fig 2A), ITS1 (Fig 2B), ITS2 (Fig 2C), 3’ ETS (Fig 2D). To ensure robustness of our data, we analyzed 3 distinct cell lines (HeLa, HAP1 and U2OS cells). As previously demonstrated, exposure to AsO_2_ results in a defect in pre-rRNA biogenesis that results in accumulation of a non-canonical 34S precursor that is emblematic of failure to process at the early A’/01 processing site but spurious processing at the “2” processing site located withing ITS1 (Fig 1). This effect was consistent in all three cell lines and resulted in significant accumulation of the 34S precursor (Fig 2E, Supp. Fig 1A). The largest product detected by 5’ETS blotting was unchanged (Fig 2E). This was confirmed by blotting with ITS1 (Fig 2B, Supp. Fig 1B) and ITS2 (Fig 2C, Supp. Fig 1C) which demonstrated slight reductions in later precursors, particularly small subunit precursors, but unaltered abundance of the largest precursors. We hypothesized that this product represented completely unprocessed 47S precursor resulting in failure to process at the immediate 5′ end (at A′/01) and a the immediate 3′ end (at 02). To determine if this were the case, we designed northern blotting probes to detect the 3’ETS containing pre-rRNAs. This would detect 47S precursors and 46S precursors that are defined by processing at A′/01 but unprocessed 3’ end. To our surprise, upon blotting using a 3′ ETS probe, we found that this largest product was significantly diminished in all three cell lines (Fig 2D, 2F, Supp. Fig 1D) and we did not find a downstream cleavage product that was complementary to the non-canonical 34S precursor found upon 5′ ETS blotting. We also find a decrease in other 3′ ETS containing mRNAs (32S-L and 28S-L). These results indicate that although primary 5’ end processing is halted by exposure to AsO_2_, primary 3′ end processing at 02 is uninhibited. Thus, contrary to our initial model, exposure to AsO_2_ does not result in accumulation of unprocessed 47S, but instead, results in accumulation of a novel precursor in which 3’ end processing precedes 5’ end processing. We term this novel intermediate the 47S-C pre-rRNA in which the “C” refers to *court* (short in French) (Fig 2G). Similar nomenclature has been ascribed to a shorten version of the 21S precursor, known as 21S-C. These findings provide the first direct evidence in human cells that 5′ and 3′ end processing of pre-rRNA can be uncoupled and proceed independently. Notably, we show that 3′ end cleavage at site 02 can occur prior to 5′ end processing at A′/01, representing the first such observation in mammalian cells. This reveals previously unrecognized flexibility in the earliest steps of rRNA maturation.

**Figure 2.**
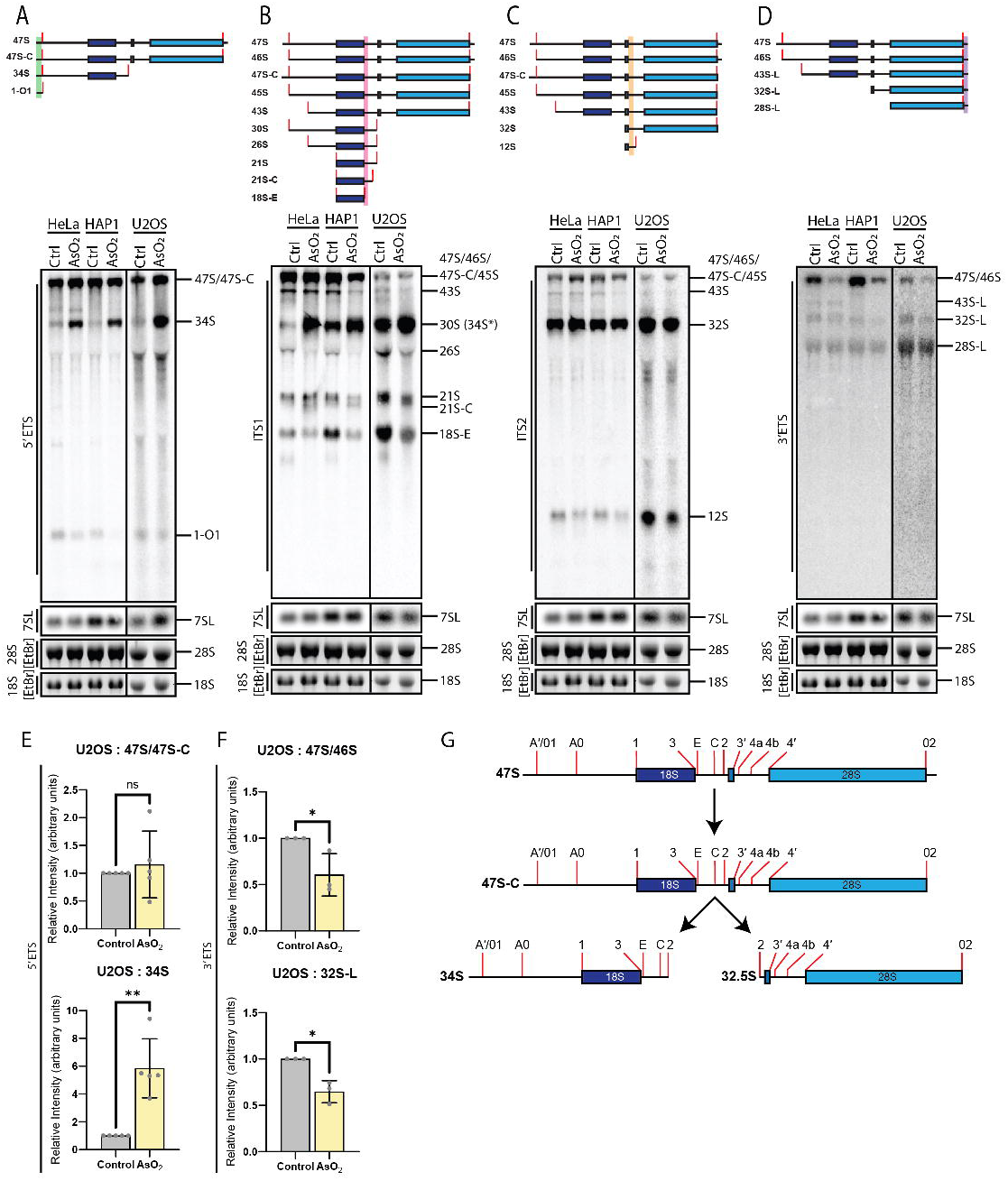
Northern blotting of HeLa, HAP1, and U2OS cell lines treated with 250 μM AsO_2_ for 2 hrs shows decoupling of 47S 5’ and 3’ end processing. (**A**) Northern blot of treated cell lines with 5’ETS probe. Treatment with AsO_2_ results in lack of processing for 47S/novel 47S-C precursors and accumulation of non-canonical 34S precursor. Blot of 7SL probe and 28S/18S ethidium bromide (EtBr) images included for loading. Simplified graphical representation of precursor species are shown above each representative blot (n=3). (**B**) ITS1 Northern probe shows selective clearance (47S/46S/45S, 43S, 26S, 21S, 21S-C, 18S-E) or accumulation (30S(34S*)) of key precursors upon sodium arsenite stress. (**C**) Processing of precursors captured with ITS2 Northern blotting is largely unperturbed upon sodium arsenite treatment. (**D**) 3’ETS Northern blot probe shows clearance of largest precursors (47S/46S) and subsequent clearance of all downstream precursors (43S-L, 32S-L, 28S-L). (**E**) Quantification of 5’ETS Northern blot U2OS 5’ETS 47S/novel 47S-C and 34S species (n=4) shows significant 34S increase upon AsO_2_ treatment in U2OS cells, analyzed by t test (*ns P > 0.05, ** P < 0.01*). (**F**) Quantification of large precursors from U2OS 3’ETS blotting (47S/46S) and 32S-L (n-3) significantly decreases upon sodium arsenite treatment *(* P < 0.05).* (**G**) Simplified processing diagram of quantified precursor species.

### Endoplasmic Reticulum stress does not phenocopy AsO_2_-induced stalling of pre-rRNA processing

AsO_2_ is a potent activator of the ISR, a pathway that inhibits protein synthesis via phosphorylation of eIF2α. AsO_2_ activates heme-regulated inhibitor (HRI) to phosphorylate eIF2α. We have previously demonstrated that stalling at the A′/01 processing site occurs concurrently with eIF2α phosphorylation, but is not dependent upon it (2). The stalling is induced not only by AsO_2_, but also by lomustine, a DNA alkylating agent used in glioblastoma treatment. Notably, we recently showed that lomustine also activates the ISR through HRI, similar to AsO_2_ (28). These findings suggest that A′/01 stalling may be a common feature of HRI-mediated ISR activation, although not directly driven by eIF2α phosphorylation. To test whether A′/01 processing defects are generally correlated to ISR activation or reflect a distinct stress response, we examined the effects of endoplasmic reticulum (ER) stressors that activate the ISR via PERK, a different eIF2α kinase. We treated with tunicamycin (Tun), an inhibitor of N-linked glycosylation; thapsigargin (Tg), which dysregulates Ca^2+^ flux in the ER, and dithiothreitol (DTT), a reducing agent that disrupts ER-mediated protein folding. Surprisingly, exposure to ER stressors had minimal effects on rRNA biogenesis (Figure 3, Supp. Fig 2-3). In contrast to AsO_2_, we did not observe any alteration in the abundance of the 47S or 47S-C species (Fig 3A, 3B), nor did we detect accumulation of the non-canonical 34S precursor (Fig 3A, 3C) following exposure to any ER stressor at any of the three doses tested. We did observe a modest but statistically significant increase in 30S levels (Fig 3A, 3D), suggesting a potential disruption in late-state small subunit maturation that warrants further investigation. No significant accumulation of the canonical 32S intermediate was detected (Fig 3A, 3E).

**Figure 3.**
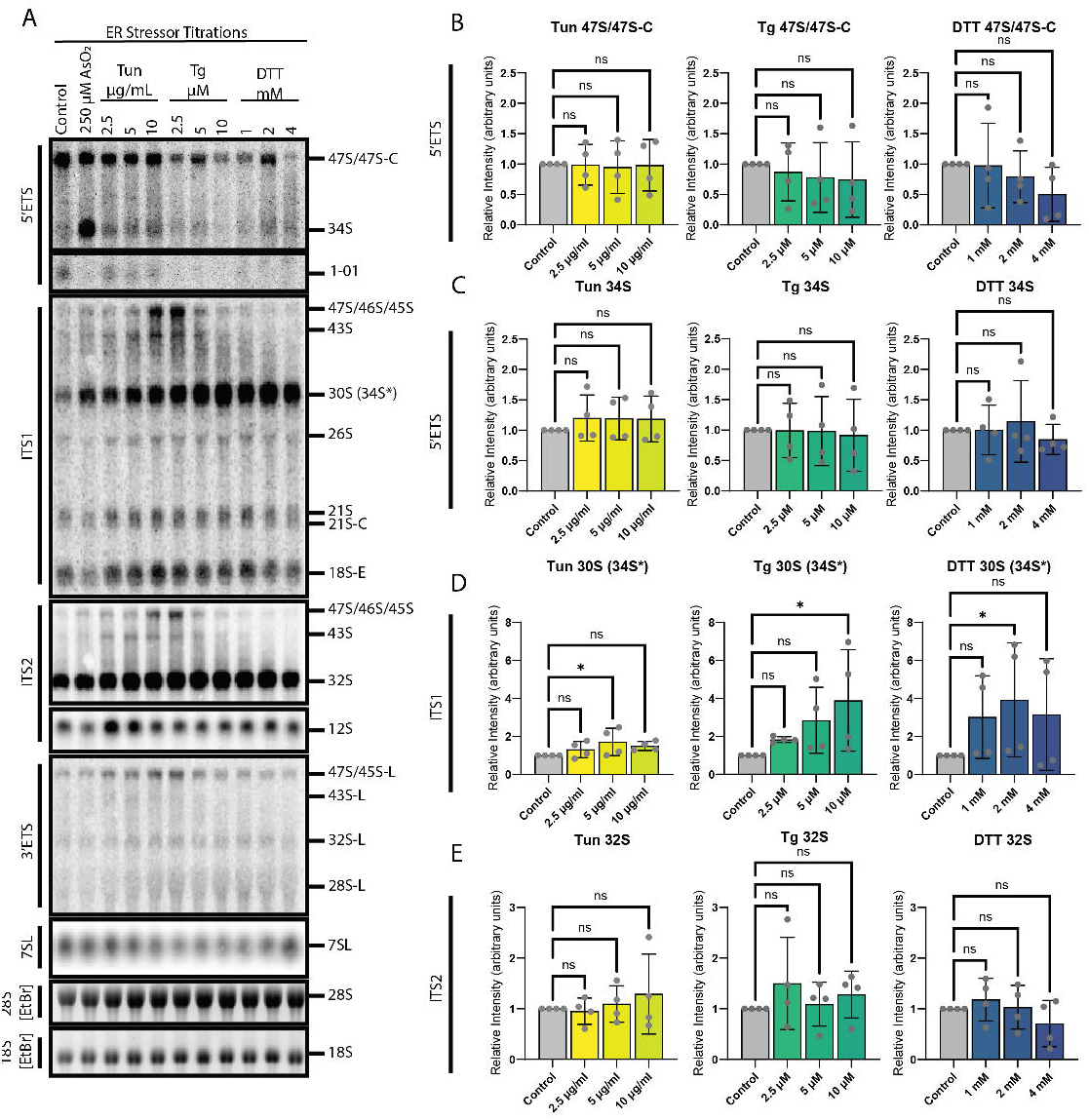
Endoplasmic reticulum (ER) stressors do not cause significant perturbations in pre-rRNA processing. (**A**) RNA from U2OS cells treated with 250 μM AsO_2_ and titrations of Tunicamycin (Tun) (2.5, 5, 10 μg/mL), Thapsigargin (Tg) (2.5, 5, 10 μM), and dithiothreitol (DTT) (1, 2, 4 mM) run on a Northern blot and probed with a panel of probes mapped to transcribed spacer regions (5’ETS, ITS1, ITS2, 3’ETS), and 7SL. Ethidium bromide (EtBr) images of mature 28S and 18S rRNA included for loading. Sodium arsenite significantly increases accumulation of select precursors while ER stressors have limited to no effect. (**B**) Quantification of 5’ETS 47S/novel 47S-C/46S (n=4) shows no significant change in accumulation across Tun, Tg, or DTT titrations. Analyzed by one way ANOVA with Dunnett’s multiple comparisons post-hoc test (*ns P > 0.05*). (**C**) 5’ETS 34S species quantification of ER stressor titrations show no significant change compared to control. (**D**) Modest increases in accumulation of ITS1 30S(34S*) precursor in Tun, Tg, and DTT titrations (*\* P < 0.05, ** P < 0.01*). (**E**) ITS2 32S quantification shows no significant change in species accumulation across Tun, Tg, and DTT ER stressor titrations.

These findings contradict our prior model in which A′/01 stalling was thought to occur in parallel with ISR activation. To further test whether A′/01 stalling is specifically linked to HRI-mediated stress rather than ISR activation more broadly, we confirmed that each ER stressor effectively inhibited global protein synthesis as expected.

Through ^35^S-metabolic labeling, we observed a dose-dependent decrease in global protein synthesis following treatment with all three ER stressors as expected (Fig 4A, 4B). This correlated with a dose-dependent increase in eIF2α phosphorylation (Fig 4C, 4D), confirming that each compound effectively activates the ISR. However, despite ISR activation, none of the ER stressors triggered A’/01 stalling or altered pre-rRNA processing (see Figure 3). The extent of eIF2α phosphorylation differed across treatments with tunicamycin inducing a modest but statistically significant increase, whereas thapsigargin and DTT triggered robust phosphorylation. Importantly, none of the ER stressors inhibited mTOR signaling, as indicated by sustained phosphorylation of 4EBP (Fig 4C, 4E). Given that rRNA is sensitive to mTOR-dependent signaling (16,29), this may explain the persistence of pre-rRNA species under ER stress conditions.

**Figure 4.**
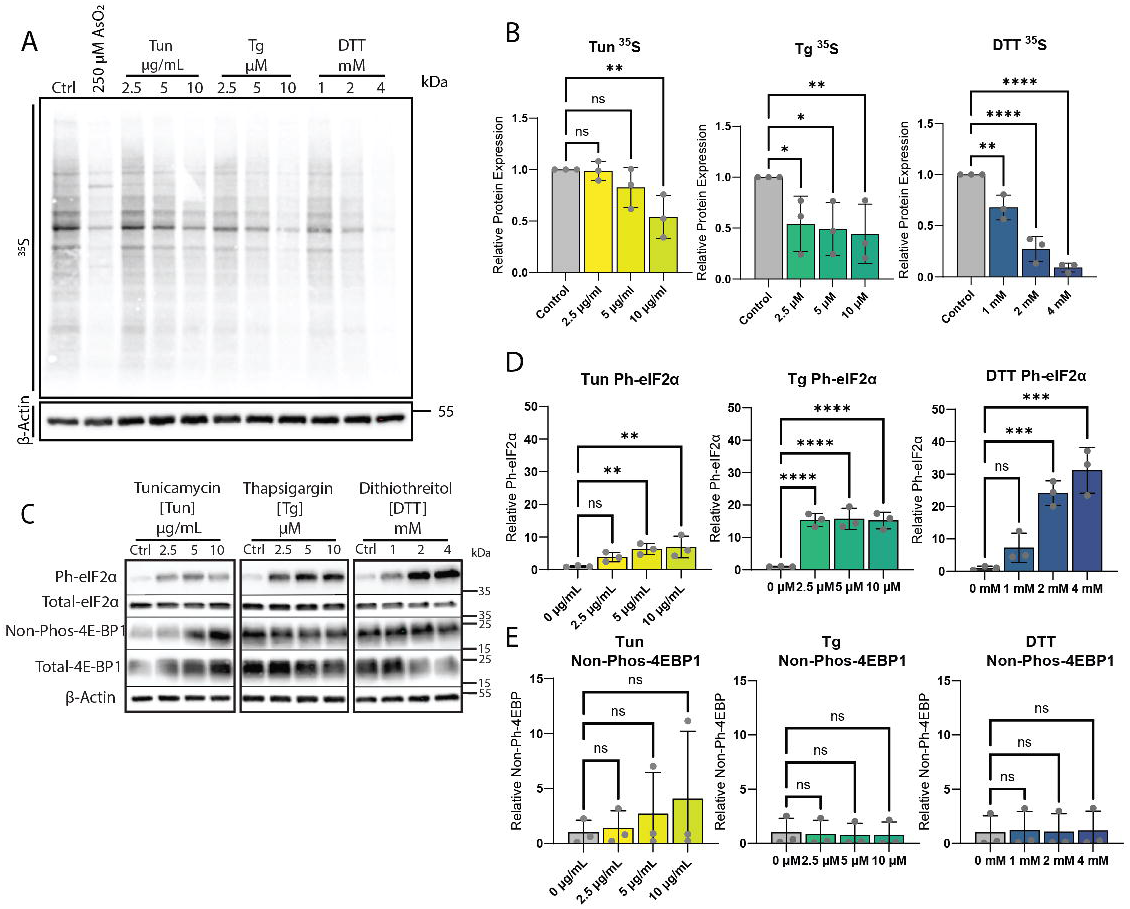
ER stressors downregulate translation and activate integrated stress response (ISR) signaling in a dose dependent manner, and do not inactivate mammalian target of rapamycin (mTOR) signaling. (**A**) ^35^S metabolic labeling of U2OS cells treated with 250 μM AsO_2_ and titrations of Tun, Tg, and DTT show effect on downregulating translation. Western blot of β-Actin shown for loading of metabolic labeling blot. (**B**) Quantification of relative protein expression of ^35^S metabolic labeling blot (n=3) shows dose dependent decrease in translation for Tun, Tg, and DTT, with DTT showing the most dramatic effect. Data analyzed by one way ANOVA with Dunnett’s multiple comparisons post-hoc test (*ns P > 0.05*, *\* P < 0.05, ** P < 0.01, *** P < 0.001, **** P < 0.0001*). (**C**) Western blotting of Ph-eIF2α, total-eIF2α, non-Phos-4EBP1, total-4EBP1, and β-Actin show ER stressors dose dependently increase phosphorylation of eIF2α, and do not dephosphorylate 4EBP1. (**D**) Quantification of Western blots show Tun, Tg, and DTT significantly phosphorylate eIF2α in a dose dependent manner, with DTT having the most profound effect. (**E**) Quantification of non-Phos-4EBP1 blots show ER stressors have no significant effect on dephosphorylation of 4EBP1 at any of the tested concentrations.

### ER-stressors decouple alterations in nucleolar morphology and stress granule formation

Our previous work demonstrated that AsO₂-induced stalling at the A’/01 processing site leads to increased nucleolar circularity, coinciding with cytoplasmic stress granule formation (2). The nucleolus, a prototypical LLPS organelle, exhibits fluid-like behavior: when pre-rRNA synthesis is suppressed, it minimized surface tension by adopting a more spherical morphology (5,30). Consistent with our observation that pre-rRNA processing remains intact under ER stress, we detected no significant changes in nucleolar morphology across all ER stressors and concentrations, as assessed using BMS1, a marker of the dense fibrillar component where early rRNA processing occurs (Fig 5A, 5B). To further assess nucleolar function, we evaluated *de novo* pre-rRNA synthesis using 5-ethynyl uridine (5EU) labeling. None of the ER stressors abolished nucleolar RNA synthesis (Fig 5C, 5D), despite clearly inhibiting global protein synthesis (Fig 4A, 4B). Thapsigargin treatment did cause a slight but significant decrease in 5EU at the highest concentration (Fig 5D), suggesting a partial suppression of transcription.

**Figure 5.**
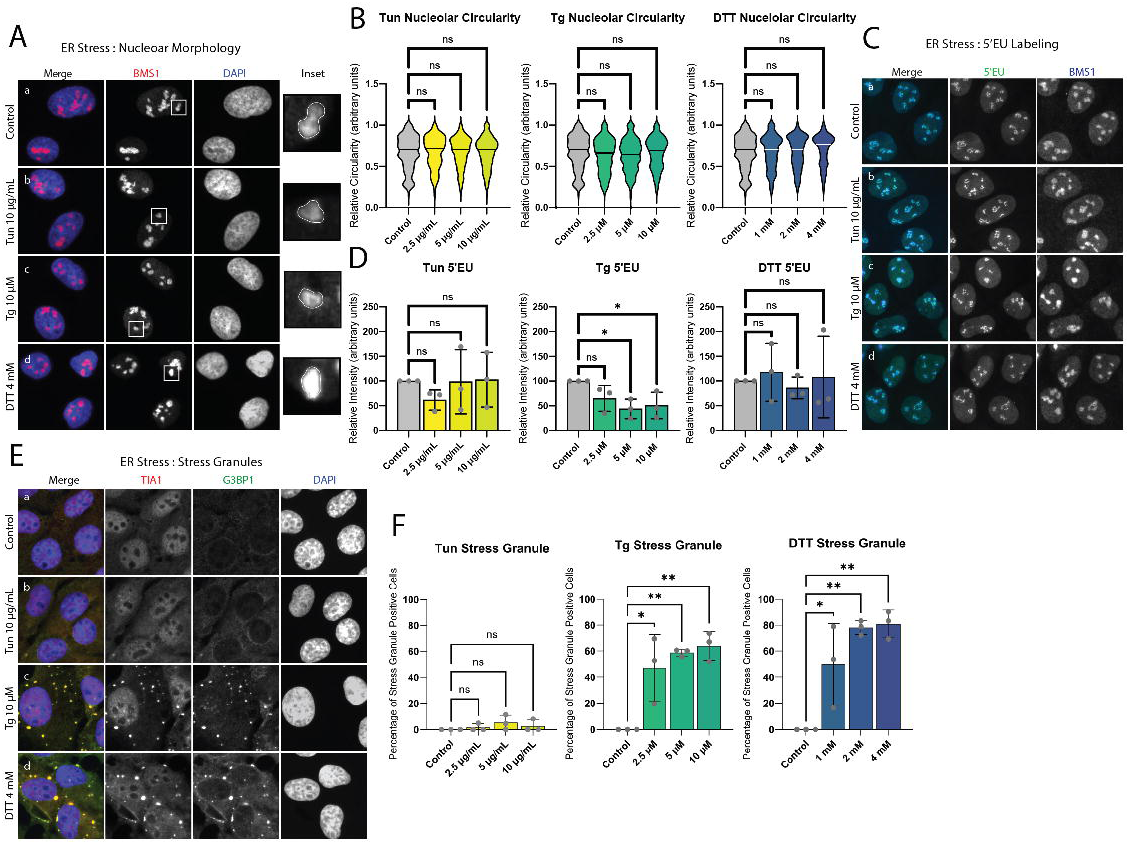
ER stressors do not modulate nucleolar morphology, have limited effect on active cellular transcription, and some cause the formation of stress granules at the tested concentrations. (**A**) Immunofluorescence (IF) of U2OS cells untreated (Aa) or treated with Tun 10 μg/mL (Ab), Tg 10 μM (Ac), and DTT 4 mM (Ad) stained with BMS1 (red) to monitor nucleolar morphology (n=3 images per condition from 3 biological replicates). (**B**) Nucleolar morphology described by scoring relative circularity of nucleoli is unchanged under all titration concentrations of applied ER stressors. Mixed-effects analysis was used to compare data, followed by Dunnett’s multiple comparisons post-hoc test (*ns P > 0.05*). (**C**) Active cellular transcription visualized by IF using 5-ethynyl uridine (5EU) click chemistry labeling (green) of U2OS cells untreated (Ca) or treated with Tun 10 μg/mL (Cb), Tg 10 μM (Cc), and DTT 4 mM (Cd). Cells labeled with 5EU for 30 mins and co-stained with BMS1 (blue) to visualize RNA transcription within the nucleoli. (**D**) Relative intensity of 5EU quantification shows the concentrations of Tun, Tg, and DTT tested do not have a drastic effect on transcription. Tg modestly decreases transcription at higher concentrations. Data analyzed by one way ANOVA with Dunnett’s multiple comparisons post-hoc test (*ns P > 0.05*, *\* P < 0.05*). (**E**) IF of U2OS cells untreated (Ea) or treated with Tun 10 μg/mL (Eb), Tg 10 μM (Ec), and DTT 4 mM (Ed) stained with TIA1 (red) and G3BP1 (green) to visualize stress granule formation. Tg and DTT treatment robustly form stress granules at the concentrations shown, while Tun treatment does not form stress granules. (**F**) Quantification of percentage stress granule positive cells for Tun, Tg, and DTT titrations show Tg and DTT treatment dose dependently induces stress granule formation, while Tun does not (*ns P > 0.05*, *\* P < 0.05, ** P < 0.01*).

Finally, in line with ISR activation, both thapsigargin and DTT robustly induced stress granule formation at all tested doses. By contrast, tunicamycin, which elicited weaker eIF2α phosphorylation (Fig 4C, 4D), failed to trigger stress granule formation under the same conditions. These findings demonstrate that A′/01 stalling is fully decoupled from ISR activation and that changes in nucleolar morphology do not necessarily accompany stress granule formation. Moreover, the persistence of rRNA synthesis despite suppression of protein synthesis suggests that ER stress may uncouple the coordinated production of ribosomal components, potentially leading to energetic misallocation in disease states marked by chronic ER stress.

### Heavy metal stress coincides with A’/01 stalling

Given the unexpected observation that pre-rRNA processing defects and ISR activation do not always correlate, we sought to clarify how these processes are linked under conditions of heavy metal stress. To this end, we selected a panel of well-characterized environmental toxicants, including both heavy metals and metalloids, to assess whether stalling at the A′/01 cleavage site of the 47S pre-rRNA is a general feature of metal-induced stress. As a positive control, we used AsO_2_, which we previously showed disrupts early pre-rRNA processing and activates the ISR (2).

To explore mechanistically distinct forms of metal-induced stress, we next examined the effects of chromium trioxide (CrO_3_) and cadmium chloride (CdCl_2_). CrO_3_ is a hexavalent chromium [Cr(VI)] compound and a potent oxidizer that generates reactive oxygen species as Cr(VI) is reduced to Cr(V) and ultimately Cr(III) within the cell (31,32). Cr(VI) is widely accepted to be genotoxic and carcinogenic and represents a major environmental pollutant (33,34). In contrast, CdCl_2_, a divalent cadmium salt [Cd(II)], disturbs metal ion homeostasis, impairs mitochondrial function and alters transcriptional responses and is a major environmental pollutant associated with genotoxic stress and carcinogenesis (35-37). CdCl_2_ also strongly triggers a proteotoxic stress response, resulting in sharp upregulation of HSF1 and NRF2 in contrast to Cr(VI), whereas Cr(VI) does not elicit the same response (38).

Upon treatment, all metals exerted rapid and measurable effects on rRNA biogenesis at all doses (Figure 6, Supp. Fig 4-5). CdCl_2_, like AsO_2_, potently induced the formation of a non-canonical 34S precursor while having minimal impact on the abundance of the 47S and the novel 47S-C pre-rRNA (Fig 6B, 6C). In contrast, CrO_3_ exposure led to more complex effects. At low concentration, CrO_3_ promoted 34S formation, but this was accompanied by a marked reduction in 47S and 47S-C levels (Fig 6A-C). At higher concentrations, the 47S/47S-C species were further depleted and 34S accumulation diminished, suggesting that transcriptional repression becomes dominant at elevated doses.

**Figure 6.**
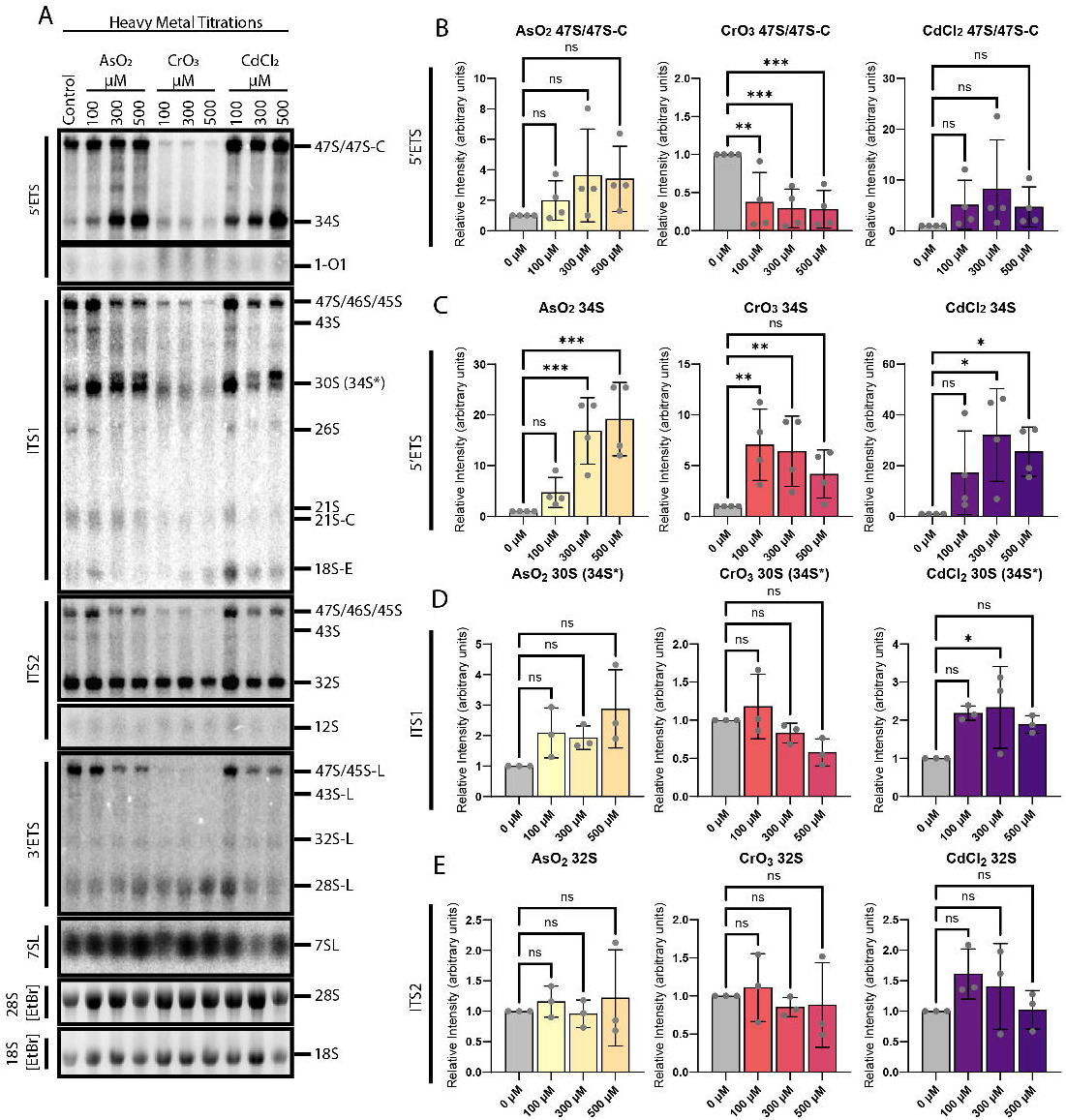
Heavy metals inhibit pre-rRNA processing at key processing checkpoints. (**A**) RNA from U2OS cells treated with titrations of AsO_2_ (100, 300, 500 μM), CrO_3_ (100, 300, 500 μM), and CdCl_2_ (100, 300, 500 μM) run on a Northern blot and probed with a panel of probes mapped to transcribed spacer regions (5’ETS, ITS1, ITS2, 3’ETS), and 7SL. Ethidium bromide (EtBr) images of mature 28S and 18S rRNA included for loading. Heavy metals have a dose dependent effect on accumulation of key precursors. (**B**) The AsO_2_ concentrations tested do not lead to significant change in accumulation of the longest pre-rRNA species (47S/novel-47S-C/46S) (n=4), while CrO_3_ significantly decreases the pool across all concentrations, and CdCl_2_ causes a modest dose dependent increase. Data analyzed by one way ANOVA with Dunnett’s multiple comparisons post-hoc test (*ns P > 0.05*, *\* P < 0.05*). (**C**) 5’ETS 34S intermediate levels dose dependently increase under AsO_2_ and CdCl_2_ treatment and are initially bolstered then diminish across the CrO_3_ titration (*\*\* P < 0.01, *** P < 0.001).* (**D**) Quantification of ITS1 30S(34S*) shows modest increase in accumulation at AsO_2_ high doses, and no significant change in levels under CdCl_2_ and CrO_3_ titrations, though CdCl_2_ trends like AsO_2_ and CrO_3_ trends in the opposite direction. (**E**) ITS2 32S quantification reveals no significant change in relative intensity of bands across all heavy metal titrations when compared to control.

These results indicate that CdCl_2_ mimics the effects of AsO_2_ on early rRNA processing, while CrO_3_ perturbs ribosome biogenesis through a distinct mechanism that involves partial A’/01 stalling but is ultimately dominated by suppression of rRNA transcription. Notably, in all conditions, intermediate precursors involved in small and large subunit maturation, specifically 30S and 32S pre-rRNAs, were retained (Fig 6A, 6D-E). These findings suggest that while early cleavage and processing steps continue under metal stress, downstream maturation events are selectively impaired. Titrations of AsO_2_, CrO_3_, and CdCl_2_ were repeated in a separate cell line to ensure robustness of the data (Supp. Fig 6). The trends of rRNA processing perturbations in HeLa cells due to heavy metal treatment largely recapitulate what was observed in U2OS cells, indicating that the processing defect is not specific to the cell line used.

### Exposure to heavy metals inhibit translation through diverse mechanisms

To better understand how protein synthesis, the ISR and A′/01 stalling are interconnected, we examined how exposure to these metals affects translation regulation. Metabolic labeling experiments reveal that all metals strongly inhibit cellular protein synthesis (Fig 7A, 7B). However, analysis of translation initiation factors uncovered distinct mechanisms that correlate with their divergent effects on rRNA processing. CdCl_2_ and AsO_2_ both triggered robust phosphorylation of eIF2α, confirming activation of the ISR (Fig 7C, 7D). While AsO_2_ signals through HRI, the upstream pathway for CdCl_2_ remains to be determined as it may involve multiple eIF2α kinases given its role in promoting ROS generation and proteotoxic stress.

**Figure 7.**
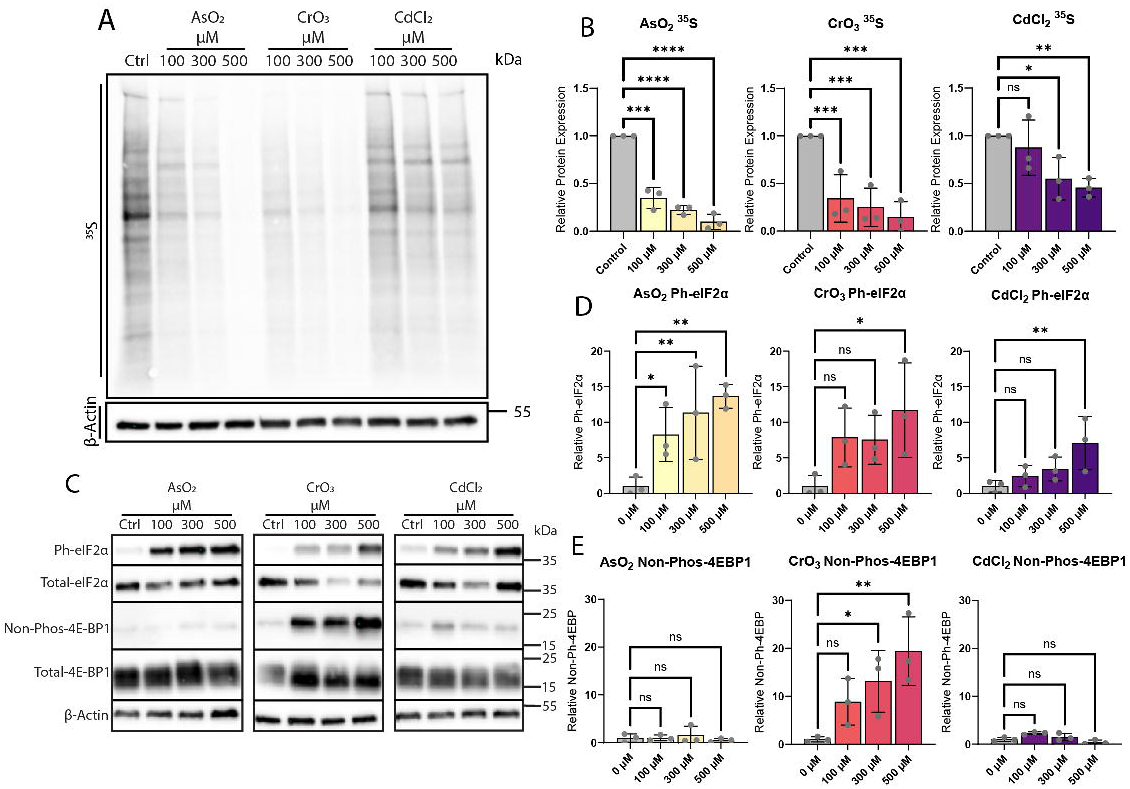
Heavy metals dramatically inhibit translation and activate ISR signaling, while CrO_3_ also inactivates mTOR signaling. (**A**) ^35^S metabolic labeling of U2OS cells treated with titrations of AsO_2_, CrO_3_, and CdCl_2_ shows strong dose dependent inhibition of active cellular translation for all heavy metals. Western blot of β-Actin shown for loading. (**B**) Quantification of metabolic labeling (n=3) showing AsO_2_, CrO_3_, and CdCl_2_ dramatically decreasing relative protein expression as concentrations increase. AsO_2_ has the most significant effect, followed by CrO_3_, and then CdCl_2_. Data analyzed by one way ANOVA with Dunnett’s multiple comparisons post-hoc test (*ns P > 0.05*, *\* P < 0.05, ** P < 0.01, *** P < 0.001, **** P < 0.0001*). (**C**) Western blotting of Ph-eIF2α, total-eIF2α, non-Phos-4EBP1, total-4EBP1, and β-Actin, show heavy metals dose dependently increase phosphorylation of eIF2α, and only CrO_3_ dephosphorylates 4EBP1. (**D**) Quantification of relative Ph-eIF2α for AsO_2_, CrO_3_, and CdCl_2_ titrations shows eIF2α is phosphorylated at the highest tested concentrations and is most robustly phosphorylated at all concentrations by AsO_2_. (**E**) Quantification of relative non-Phos-4EBP1 shows AsO_2_ and CdCl_2_ treatments do not result in dephosphorylation of 4EBP1, while CrO_3_ significantly increases dephosphorylation in a dose dependent manner.

In contrast, CrO_3_ induced strong translational repression without robust eIF2α phosphorylation, except at the highest concentrations (Fig 7A-D). Instead, CrO_3_ exposure resulted in marked dephosphorylation of 4EBP (Fig 7C, 7E), which in its hypophosporylated state binds to eIF4E and prevents formation of the eIF4F complex (39,40). Under normal conditions, 4EBP is maintained in an inactive state through phosphorylation by mTOR. mTOR inhibition therefore blocks cap-dependent translation and also reduces rRNA transcription (16,29). This mechanism of repression aligns with the observed loss of 47S/47S-C pre-rRNA following CrO_3_ treatment (Figure 6), suggesting that mTOR downregulation impairs both translation initiation and early steps in ribosome biogenesis.

Together, these findings indicate that CdCl_2_ and AsO_2_ converge on ISR-dependent pathways to inhibit translation and disrupt early rRNA processing, while CrO_3_ exerts its effects through mTOR inhibition, linking translation repression to transcription silencing of rDNA. Given the widespread environmental presence of these metals, such mechanistic differences in how translation and ribosome biogenesis are suppressed may have distinct implications for cellular adaptation, disease progression or therapeutic targeting.

We next investigated whether exposure to each metal elicited similar effects on nucleolar morphology and stress granule formation. Cells were treated with a range of concentrations for each compound and nucleolar structure was assessed by immunostaining for BMS1, as previously described (Figure 5). Unlike classical ER stress inducers, all three metals induced a pronounced change in nucleolar morphology, characterized by increased circularity and a more compact spherical appearance (Fig 8A, 8B). Notably, nucleolar integrity remained intact following CrO_3_ exposure. This was unexpected, as loss of 47S pre-rRNA, which serve as key scaffolds for nucleolar organization, typically leads to nucleolar dissolution (2,5,41). However, persistence of intermediate pre-rRNA species such as 30S and 32S, even under CrO_3_ treatment (Fig 6D, 6E), likely provides sufficient structural support to maintain nucleolar cohesion, preventing complete disassembly as observed with actinomycin D treatment (2).

**Figure 8.**
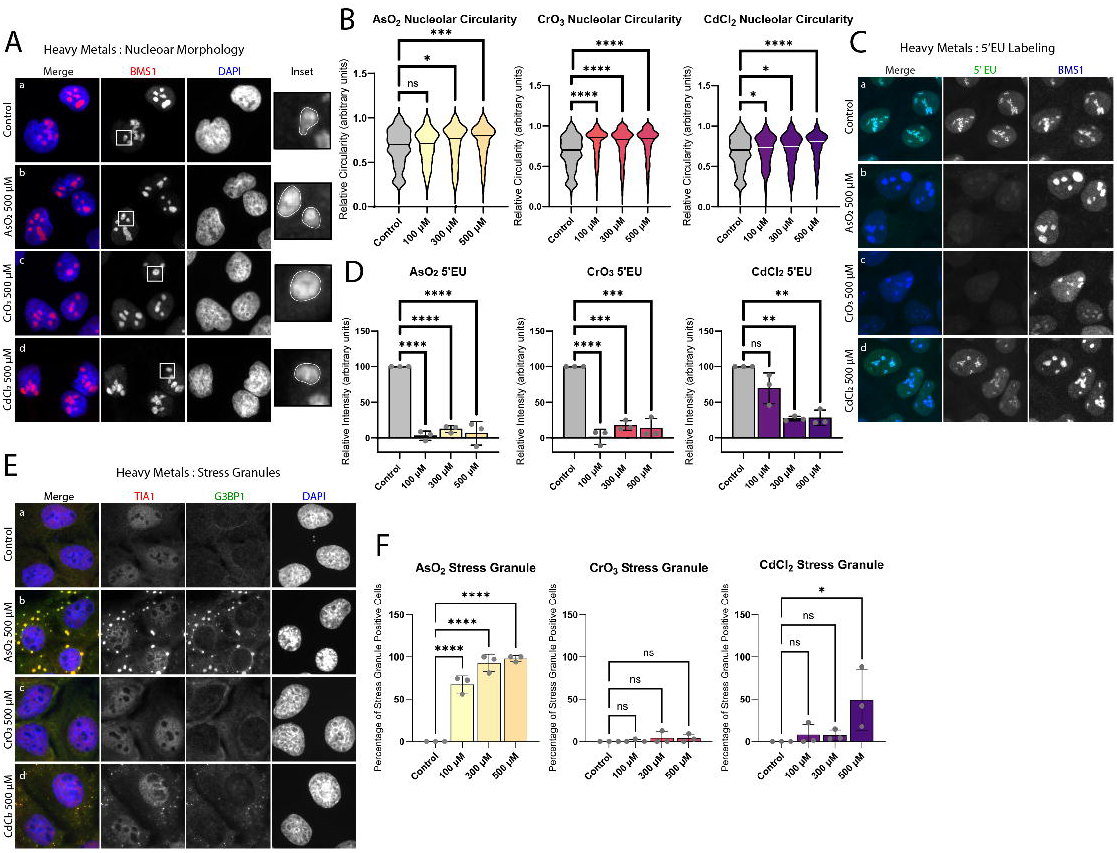
Heavy metals cause a distinct change in nucleolar morphology, decrease active cellular transcription, and can cause stress granule formation at the tested concentrations. (**A**) Immunofluorescence (IF) of U2OS cells untreated (Aa) or treated with AsO_2_ 500 μM (Ab), CrO_3_ 500 μM (Ac), and CdCl_2_ 500 μM (Ad) stained with BMS1 (red) to monitor nucleolar morphology (n=3 images per condition from 3 biological replicates). (**B**) Nucleolar morphology described by scoring relative circularity of nucleoli is significantly altered dose dependently by all heavy metals tested. Mixed-effects analysis was used to compare data, followed by Dunnett’s multiple comparisons post-hoc test (*ns P > 0.05*, *\* P < 0.05, *** P < 0.001, **** P < 0.0001*). (**C**) Active cellular transcription visualized by IF using 5-ethynyl uridine (5EU) click chemistry labeling (green) of U2OS cells untreated (Ca) or treated with AsO_2_ 500 μM (Cb), CrO_3_ 500 μM (Cc), and CdCl_2_ 500 μM (Cd). Cells labeled with 5EU for 30 mins and co-stained with BMS1 (blue) to visualize RNA transcription within the nucleoli. (**D**) Relative intensity of 5EU quantification shows all the tested heavy metals have a significant effect on active cellular transcription, with AsO_2_ and CrO_3_ almost completely ablating transcription and CdCl_2_ significantly reducing transcription at higher concentrations. Data analyzed by one way ANOVA with Dunnett’s multiple comparisons post-hoc test (*\*\* P < 0.01*). (**E**) IF of U2OS cells untreated (Ea) or treated with AsO_2_ 500 μM (Eb), CrO_3_ 500 μM (Ec), and CdCl_2_ 500 μM (Ed) stained with TIA1 (red) and G3BP1 (green) to visualize stress granule formation. AsO_2_ robustly forms classical stress granules, while CrO_3_ does not form stress granules and CdCl_2_ forms small, speckled granules. (**F**) Quantification of percentage of stress granule positive cells shows AsO_2_ treatment causes stress granule formation in almost all cells at every concentration tested. Significant stress granules were formed by the highest tested dose of CdCl_2_, though were distinct in appearance from arsenite stress granules. CrO_3_ treatment does not result in stress granule formation at any concentration.

To directly assess rRNA transcription, we performed 5EU metabolic labeling. Consistent with previous results (2), AsO_2_ completely abolished nucleolar transcription (Fig 8C, 8D). Our previous work demonstrated that A′/01 stalling precedes transcriptional inhibition, resulting in accumulation of the unprocessed rRNA which we now know to be the 47S-C pre-rRNA (Fig 2). Similarly, CrO_3_ led to a complete shutdown of rRNA transcription (Fig 8C, 8D). However, in this case, the persistence of intermediate precursors suggests that pre-rRNA processing is impaired at a later stage, rather than by A′/01 stalling. This transcriptional block is likely mediated by mTOR inactivation, as indicated by 4EBP dephosphorylation (Fig 7C, 7E). CdCl_2_ also reduced rRNA transcription, although to a lesser extent than AsO_2_ or CrO_3_, paralleling its more modest effect on global translation.

Finally, we examined stress granule formation in response to each metal. As expected, AsO_2_ strongly induced stress granules consistent with its robust activation of the ISR, phosphorylation of eIF2α and a wealth of historical data (Fig 8E, 8F). Despite causing near complete inhibition of protein synthesis, CrO_3_ failed to induce stress granules, in line with its inability to trigger eIF2α phosphorylation. CdCl_2_ induced smaller, less abundant stress granules at high concentrations, consistent with partial ISR activation.

Together, these findings demonstrate that heavy metals disrupt ribosome biogenesis and translation through distinct mechanisms. While AsO_2_ and CdCl_2_ converge on ISR-dependent pathways that inhibit translation, CrO_3_ acts through mTOR inhibition, suppressing both rRNA transcription and cap-dependent translation without activating the ISR. Importantly, we show that stalling at the A′/01 site is a stress response that can occur independently of eIF2α phosphorylation and stress granule formation, revealing previously unrecognized decoupling of key components of cellular stress response to environmental toxins.

## DISCUSSION

Here, we uncover previously unrecognized flexibility in early rRNA processing, showing for the first time that 3′ end cleavage at site 02 can precede 5′ end processing at A′/01 in human cells. This finding challenges the prevailing model that 5′ and 3′ end maturation of the 47S pre-rRNA are obligatorily coupled in humans (24,27). Indeed, under homeostatic conditions, human cells rarely exhibit detectable precursors between the 47S and 45S species. This contrasts with observations in murine cells, where 5′ end processing at the A’ site can precede 3′ end cleavage at site 6, generating a transient 46S intermediate (25-27). Processing sites A′ and 6 in mice are analogous to human A′/01 and 02, respectively.

Our findings also revise our previous model, which proposed that intact 47S pre-rRNA accumulates during A′/01 stalling (2). In retrospect, such retention of unprocessed transcripts might pose additional risks to cell viability during stress. Notably, the mechanistic details of 3′ end processing in mammalian cells remain poorly understood. Transcription termination by RNA polymerase I mediated by arrays of “Sal-Box” termination signals, which are bound by transcription termination factor 1 (TTF-I) (42,43). How termination is coordinated with cleavage at site 02 is unknown, but analogous Pol II transcriptional events may provide insight. Transcription termination and 3’ end processing of mRNAs are intrinsically linked (44). The “torpedo model” of transcription termination proposes that endonucleolytic cleavage downstream of the polyadenylation signal allows for access for exonucleolytic digestion by Rat1/Xrn2 of the downstream cleavage product (45,46). Exonucleolytic digestion allows for efficient release of RNA polymerase from the DNA template. Failure to cleave downstream of a polyadenylation signal during Pol II transcription results in transcriptional readthrough and improper termination (47,48). A similar scenario could occur at the rDNA locus: if 02 processing fails, Pol I may continue transcription into the intergenic spacer (IGS), potentially producing aberrant or excess RNA. Such spurious transcription, particularly in the absence of coordinated processing, may increase the likelihood of R-loop formation and genomic instability, posing a substantial risk to already stressed cells. Work in yeast suggests that Rat1/Xrn2, a 5’ to 3’ exonuclease is required for efficient transcription termination, which suggests that cleavage followed by a torpedo model may also take place for rRNA transcription termination (49). Recent data in yeast also demonstrates that transcription termination occurs rapidly after 3’ end cleavage (50). Further work is needed to define the precise molecular relationship between 3′ end processing, transcription termination, and nucleolar homeostasis in human cells.

A major barrier to understanding early pre-rRNA processing in humans is that the enzymes responsible for cleavage at A′/01 and 02 remain unidentified. Much of what is known about rRNA processing derives from *Saccharomyces cerevisiae*, a genetically tractable model organism that has yielded critical insights into ribosome biogenesis. However, the regulatory events we describe here appear to be not conserved in fungi. Notably, *S. cerevisiae* lacks a 5′ end processing site analogous to A′/01, rendering the system unsuitable for identifying the corresponding endonuclease in humans. However, it is established that this early cleavage event in mammals depends on the U3 snoRNP, which is required for proper formation of the small subunit processome (51). Similarly, while 3′ end processing in yeast occurs downstream of the 25S rRNA at site B0, this reaction is mediated by the RNase III enzyme Rnt1p, followed by exonucleolytic trimming by Rex1p (52,53). The mammalian homolog of Rex1p, known as REXO5/NEF-sp, is largely restricted to the germline, and its deletion in mice results in viable fertile offspring, suggesting it is unlikely to play a general role in somatic ribosome biogenesis (54). As with A′/01 processing, 3′ end cleavage in mammals also appears to require a snoRNP, in this case, the U8 snoRNP (55,56). These differences underscore the need for mammalian-specific models to dissect early processing events and identify the endonucleases involved.

Our results also reveal how distinct forms of stress can differentially affect the coordination between ribosome biogenesis, protein synthesis, and rRNA transcription. During cellular stress, anabolic pathways are generally suppressed to conserve energy. Given the substantial energetic cost of ribosome production, it is unsurprising that this process is subject to multiple regulatory checkpoints. A particularly unexpected finding from our study is that activation of ER stress pathways has minimal impact on rRNA transcription, despite causing a sharp reduction in global protein synthesis.

mRNAs encoding ribosomal proteins are especially sensitive to translational repression due to the presence of a 5′ terminal oligopyrimidine (TOP) motif, which confers tight translation regulation in response to stress (57-59). As a result, continued rRNA synthesis during ER stress may lead to an imbalance in ribosomal RNA and ribosomal protein production. This asymmetry could contribute to proteostatic stress, inefficient ribosome assembly, and energetic waste. In neurodegenerative disorders marked by chronic ER stress (60-62), such as amyotrophic lateral sclerosis (ALS), Parkinson’s disease, and Alzheimer’s disease, this imbalance may contribute to disease pathogenesis. Notably, small-molecule inhibitors of PERK or ISR signaling, which restore translation during chronic ER stress, have shown promise in models of prion disease, Parkinson’s disease and frontotemporal dementia (63-67). Importantly, these interventions do not resolve the underlying ER stress but instead permit continued translation despite ongoing cellular stress. Our findings suggest that such compounds may help re-establish the balance between rRNA synthesis and ribosomal protein production, potentially mitigating ribosome assembly defects. However, further investigation is needed to determine whether this restored translational capacity results in fully functional ribosomes. Future studies should examine how chronic ER stress affects rRNA processing and assess the integrity and translational fidelity of ribosomes produced under these conditions.

Finally, our results identify metal-induced stress as a potent and specific trigger of A′/01 stalling (Figure 9). By examining a panel of environmentally and clinically relevant metals, we uncovered distinct mechanisms by which these agents disrupt rRNA processing and translation regulation. Both AsO_2_ and CdCl_2_ robustly induced A′/01 stalling and accumulation of the non-canonical 34S precursor, in parallel with strong phosphorylation of eIF2α and inhibition of protein synthesis, consistent with activation of the ISR. In contrast, CrO_3_ disrupted ribosome biogenesis through a distinct pathway: despite failing to activate the ISR or induce stress granules, CrO₃ exposure led to marked dephosphorylation of 4EBP, suppression of mTOR signaling, and loss of 47S/47S-C pre-rRNA species. CrO_3_ also resulted in a modest increase in non-canonical 34S; however, the generation of this product was blunted as 47S/47S-C levels decreased as these are the direct precursors for 34S production.

**Figure 9.**
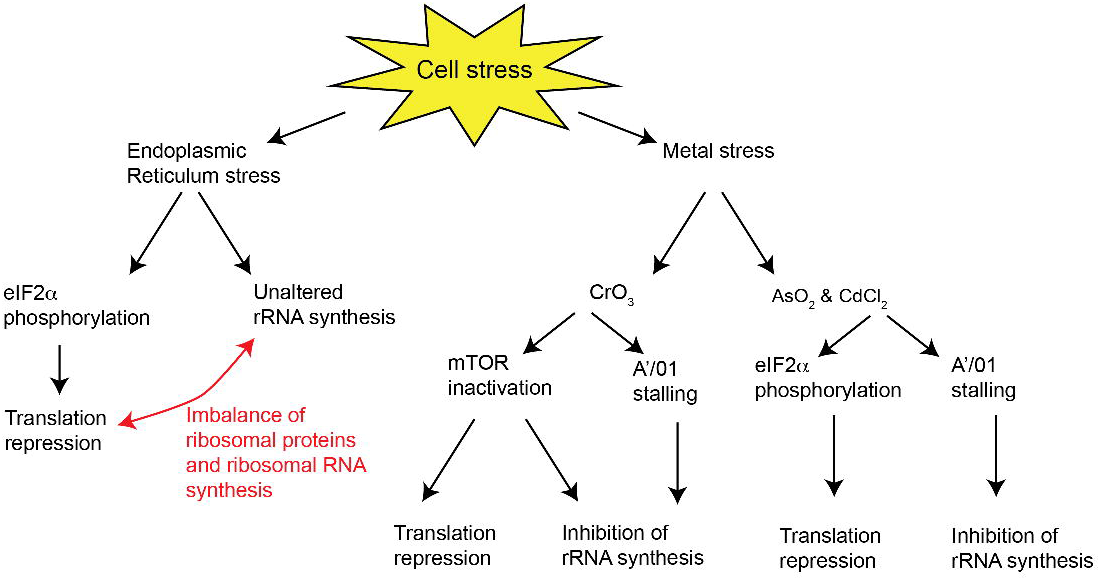
Model of heavy metal and ER stressors differential effects on distinct stress pathways. ER stressors cause phosphorylation of eIF2α and translation repression while not effecting rRNA synthesis. This creates an imbalance of ribosomal proteins and ribosomal RNA. Outcomes of heavy metal stress are distinct between CrO_3_ and AsO_2_ & CdCl_2_. Chromium stress causes mTOR inactivation and A’/01 stalling which leads to translational repression and inhibition of rRNA synthesis in tandem. Arsenite and cadmium alternatively cause phosphorylation of eIF2α and A’/01 stalling without effecting mTOR, and lead to translational repression and inhibition of rRNA synthesis.

Arsenite, cadmium, and chromium all generate ROS and are genotoxic to different extents. CrO_3_ exerts effect through direct DNA damage and is localized mainly to the nucleus, causing double and single strand breaks as well as forming Cr-DNA adducts (38,68). AsO_2_ and CdCl_2_ alternatively are localized to the cytoplasm and act through the generation of ROS both directly and indirectly (38). Difference in localization between AsO_2_ and CdCl_2_ compared to CrO_3_ indicate potential reason for differences in the stress responsive pathways we show to be activated by the respective metals.

CrO_3_-induced stress also reveals novel aspects of nucleolar biology. Historically, active rDNA transcription has been considered essential for maintaining nucleolar integrity, as classic transcriptional inhibitors like actinomycin D cause rapid nucleolar dissolution (69,70). Our earlier discovery of A′/01 stalling showed that nucleolar structure could be preserved even when transcription was impaired, provided the largest precursor, now identified as 47S-C, was retained (2). In contrast, our current analysis demonstrates that even in the absence of all large precursors (47S, 47S-C, 46S, and 45S) and active rRNA transcription, nucleolar integrity can be maintained if intermediate precursors persist. Following CrO_3_ exposure, we observed depletion of these early precursors while the nucleolus remained intact, suggesting that species such as 30S and 32S pre-rRNAs are sufficient to sustain nucleolar organization. These findings indicate that nucleolar integrity depends not only on active transcription but also on the presence of a critical pool of pre-rRNA intermediates that maintain phase separation and structural cohesion.

These findings highlight that A′/01 stalling is not a universal feature of translation inhibition or ISR activation, but rather a stress-specific response predominantly triggered by certain heavy metals. This specificity suggests that metal-induced cellular damage engages unique signaling or structural checkpoints in rRNA processing that are distinct from canonical stress responses such as ER stress. Identifying the molecular mediators of these metal-specific effects will be critical for understanding how environmental toxins reshape nucleolar function and ribosome biogenesis. It is worth pointing out that while metal stress appears particularly potent in triggering A′/01 stalling, this phenomenon is not restricted to metal exposure. Our initial work revealed that the chemotherapeutic agent lomustine also induces A′/01 stalling, likely through HRI-dependent ISR activation (2,28). Together, these findings suggest that A′/01 stalling may serve as a convergent point for diverse stress pathways that impinge on nucleolar function. Future work aimed at defining the upstream sensors, endonucleases, and RNA-binding factors involved in this response will be essential for fully understanding how cells regulate ribosome biogenesis under adverse conditions and how this regulation is rewired in disease states.

Together, our findings reveal that early rRNA processing is more flexible than previously appreciated and can be selectively disrupted by distinct forms of cellular stress. We identify A′/01 stalling as a stress-specific regulatory event that can be triggered independently of ISR activation and stress granule formation, with heavy metals representing particularly potent inducers. These results uncover a previously unrecognized layer of regulation in ribosome biogenesis and highlight the nucleolus as an active sensor of environmental and therapeutic insults. Understanding how cells modulate rRNA processing in response to stress will be essential for uncovering new principles of stress adaptation and may offer novel entry points for therapeutic intervention in diseases marked by translational imbalance or nucleolar dysfunction.

## METHODS

### Antibodies

Total-eIF2α (Santa Cruz, sc-133132), phospho-eIF2α (AbCam, ab32157), total-4E-BP1 (Cell Signaling Technology, 9452), non-phospho-4E-BP1 (Cell Signaling Technology, 4923), β-Actin (Cell Signaling Technology, 3700), BMS1 (Santa Cruz, sc-271040), TIA1 (Proteintech, 12133-2-AP), G3BP1 (Santa Cruz, sc-365338), Cy2 Goat anti-mouse IgG, FcL subclass 1 (Jackson ImmunoResearch, 115-225-205), Cy3 Goat anti-mouse IgG, FcL subclass 1 (Jackson ImmunoResearch, 115-165-205), Cy3 Goat anti-mouse IgG, FcL subclass 2b (Jackson ImmunoResearch, 115-165-207), Cy5 Goat anti-mouse IgG, FcL subclass 2b (Jackson ImmunoResearch, 115-175-207), Cy5 xGoat anti-Rabbit IgG (H+L) (Jackson ImmunoResearch, 111-175-144), Affipure Goat Anti-Mouse IgG, FcL subclass 1 specific (Jackson ImmunoResearch, 115-035-205), Peroxidase AffiniPure Goat Anti-Mouse IgG, Fcy subclass 2b Specific (Jackson ImmunoResearch, 115-035-207), Peroxidase AffiniPure Goat Anti-Rabbit IgG (H+L) (Jacson ImmunoResearch, 111-035-144).

### Cell culture and drug treatment

U2OS, HAP1, and HeLa cells were maintained in DMEM (Gibco) supplemented with 10% fetal bovine serum (Atlas Biologicals) and 1% Penicillin/Streptomycin (Corning) in a 37°C incubator at 5% CO_2_. Sodium (meta)arsenite, NaAsO_2_, 98% (Sigma), Chromium(VI) oxide, CrO_3_, 99% (Thermo Scientific Chemicals), Cadmium chloride hydrate, CdCl_2_, 99.99% (metals basis) (Thermo Scientific Chemicals), Tunicamycin Mixture (Cayman Chemical), Thapsigargin (Cayman Chemical), Dithiothreitol, DTT (Fisher BioReagents) were added for 2 hrs at the concentrations indicated.

### Epifluorescence immunofluorescence

Cells were grown to ∼80% confluence on glass coverslips, stressed with indicated stressors for 2 hrs, then fixed with 4% paraformaldehyde in PBS for 15 mins and permeabilized in 0.5% Triton X-100 in PBS for 10 mins. Fixed and permeabilized cells were blocked in 5% horse serum/1X TBS in PBS for 1 hr. Cells were then incubated rocking at 4°C overnight in primary antibody and subsequently incubated for 1 hr rocking at room temperature in secondary antibody. Primary antibodies were suspended in blocking buffer with the addition of 0.2% sodium azide, secondary antibodies were suspended in blocking buffer. Secondary antibodies (Jackson ImmunoResearch) were tagged with either Cy2, Cy3, or Cy5. Cells were washed with PBS and mounted to glass slides in polyvinyl mounting media. Slides were viewed at room temperature on a Zeiss Axio Observer Z1 microscope equipped with a digital camera (C10600/ORCA-R2 Hamamatsu Photonics). Images were taken using Zen software and compiled using ImageJ Plugin QuickFigures and Adobe Illustrator 2023. Nucleolar morphology was segmented using ilastik and analyzed using a custom Fiji script. Ilastik was employed to precisely define the outline of nucleoli within the set background of DAPI nuclear signal. The ilastik derived mask for the nucleoli was then used to threshold microscopy images in ImageJ and circularity was quantified. The circularity metric in ImageJ measures circularity corrected for aspect ratio. Three sets of images per condition from 3 biological replicates were analyzed in this manner.

### 5’EU Metabolic labeling

Cells were grown and treated as noted for immunofluorescence. After drug treatment, cells were treated with 1 mM 5-ethynyl uridine labeling reagent (Click Chemistry Tools) for 30 mins. Cells were then fixed and permeabilized as noted for immunofluorescence and detect reagent – Tris [pH 8.5], 1 mM CuSO_4_, 25 uM Azide 488 (Click Chemistry Tools), 100 mM Ascorbic Acid – was added for 30 mins. Further blocking, primary, and secondary incubation were carried out as described for immunofluorescence. Relative intensity of signal from 5’EU (green) channel was used to quantify nascent transcription in ImageJ. Mean intensity of background was subtracted and average intensity across 3 sets of images per condition from 3 biological replicates was captured by an automated macro script for quantification.

### Northern blotting

Cells were grown to ∼80% confluency and treated as specified for 2 hrs. Cells were lysed using 0.8 M Guanidinium thiocynate, 0.4 M Ammonium thiocynate, 0.1 M Sodium Acetate [pH 5.0], 5% glycerol, 48% Phenol. RNA was extracted by addition of 0.5 volumes of 100% chloroform followed by centrifugation and precipitation of the aqueous phase with 2 volumes of 100% ethanol. 5-10 ug of RNA was run on a 1.2% Agarose gel, with 1X H-E buffer (20 mM HEPES, 1 mM EDTA [pH 7.8]) and 7% formaldehyde, in 1X H-E buffer at room temperature for 18 hrs at 55 V with free buffer exchange. The RNA was transferred to Amersham Hybond - N+ membrane (Cytiva) via passive evaporation and UV crosslinked at 260 nM with 1200 μJoules (X100) twice in a Stratalinker. The membranes were then stained with methylene blue and prehybridized for 1 hr at 65°C with 10 mL of ULTRAhyb Ultrasensitive Hybridization Buffer (Invitrogen). The hybridization buffer was then replaced with 10 mL of fresh buffer and allowed to reach temperature of 65°C. Pre-prepared ^32^P end labeled radioactive probe in 10 uL aliquots were denatured at 95°C for 10 mins. 1 mL of pre-warmed hybridization buffer was added to the 10 uL of denatured probe and then added back to the hybridization tube. The probe was incubated for 1 hour at 65°C then the temperature was reduced to 37°C overnight. Blots were then washed with 2X SSC / 0.1% SDS two times for 15 mins at 42°C, wrapped in cling wrap, and exposed on a Molecular Dynamics Kodak Storage Phosphor Screen S0230 overnight. Storage screens were then imaged on a 2020 Amersham Typhoon Phosphor Imager (GE), images were processed and quantified with ImageJ or Image Lab (BioRad) and compiled with Adobe Illustrator 2023. Imaged blots were stripped and prepared for re-probing by pouring around 100 mL of boiling 0.1X SSC / 0.05% SDS and rocking on a shaker until reaching room temperature three times.

### Northern Probe Preparation

Northern probes were designed using UCSC Genome Browser. DNA oligonucleotides were synthesized by Integrated DNA Technologies and resuspended in diH_2_O to a final concentration of 6 uM. To end label the DNA, 1 uL of 6 uM DNA oligonucleotide, 14 uL diH_2_O, 2 uL 10X T4 PNK buffer, 1 uL T4 PNK (NEB), and 5 uL of [γ-^32^P] ATP (3000 Ci/ml) (Perkin-Elmer) were incubated at 37°C for 1 hr. Post incubation, 80 uL of diH_2_O was added to the reaction mix and unincorporated radioactive nucleotides were filtered from the final probe pool by gel filtration in a G-25 spin column (Cytiva) according to manufacturer’s protocols. Sequence of the probes used in Northern blotting are noted here: Human 5’ETS (CGGAGGCCCAACCTCTCCGACGACAGGTCGCCAGAGGACAGCGTGTCAGC) Human ITS1 (CCTCGCCCTCCGGGCTCCGGGCTCCGTTAATGATC) Human ITS2 (CTGCGAGGGAACCCCCAGCCGCGCA) Human 3’ETS (GGAAGGACGGACGGCGCCGAACGCG) Human 7SL (GCTCCGTTTCCGACCTGGGCC).

### Western Blotting

Cells were grown to ∼80% confluence and dosed with indicated treatments for 2 hrs. Cells were then lysed with NP-40 lysis buffer – 20 mM Tris (Fisher BioReagents) [pH 8.0], 150 mM NaCl (Fisher BioReagents), 0.5% NP-40 alternative (Millipore) – and run on a 12% polyacrylamide stain-free gel (BioRad) with a 1X sodium dodecyl sulfate (SDS) running buffer for 35 mins at 240 V. Gels were imaged on a ChemiDoc Imaging System (BioRad) and transferred via Trans-Blot Turbo Transfer System (BioRad). Blots were blocked in a 5% milk/1X TBST solution rocking at room temperature for 1 hr, then washed with 1X TBST and probed overnight at 4°C with primary antibody in 1X TBS/5% Normal Horse Serum (NHS)/0.2% sodium azide (Fisher BioReagents). Blots were then washed and probed with secondary HRP antibody (BioRad) rocking at room temperature for 1 hr and imaged using Clarity ECL reagent (BioRad) on a ChemiDoc Imaging System (BioRad). Images were quantified using Image Lab software (BioRad) and compiled in Adobe Illustrator 2023.

### 35S Metabolic Labeling

Cells were grown to ∼80% confluence and dosed with the indicated treatments for 2 hrs. Cells were then placed in DMEM, high glucose, no glutamine, no methionine, no cystine media (Gibco) for 15 mins, then washed with PBS and placed in complete DMEM media with EasyTag EXPRESS^35^S Protein Labeling Mix (Perkin-Elmer) to a final concentration of 1 uCi/mL for 15 mins. Cells were harvested, lysed, run, imaged, and transferred as described in the Western Blotting segment. Following transfer, the blot was then dried at room temperature and placed directly on a Molecular Dynamics Kodak Storage Phosphor Screen S0230 overnight. Storage screens were then imaged on a 2020 Amersham Typhoon Phosphor Imager (GE), images were processed and quantified with ImageJ and compiled with Adobe Illustrator 2023. Blots were then rehydrated and blocked and probed as described in the Western Blotting segment.

### Statistics

Statistical tests and graphical representation achieved in GraphPad Prism. Mixed-effects analysis, ANOVA, and paired t tests were used where appropriate. Dunnett’s multiple comparisons test was used where appropriate for post-hoc analysis (*\* P < 0.05, ** P < 0.01, *** P < 0.001, **** P < 0.0001*).

## REFERENCES

1. Warner, J.R., Vilardell, J. and Sohn, J.H. (2001) Economics of ribosome biosynthesis. Cold Spring Harb Symp Quant Biol, 66, 567–574.

2. Szaflarski, W., Lesniczak-Staszak, M., Sowinski, M., Ojha, S., Aulas, A., Dave, D., Malla, S., Anderson, P., Ivanov, P. and Lyons, S.M. (2022) Early rRNA processing is a stress-dependent regulatory event whose inhibition maintains nucleolar integrity. Nucleic Acids Res., 50, 1033–1051.

3. Ojha, S., Malla, S. and Lyons, S.M. (2020) snoRNPs: Functions in Ribosome Biogenesis. Biomolecules, 10.

4. Johnston, R., Aldrich, A. and Lyons, S.M. (2024) Roles of ribosomal RNA in health and disease. Frontiers in RNA Research, 1.

5. Lafontaine, D.L.J., Riback, J.A., Bascetin, R. and Brangwynne, C.P. (2021) The nucleolus as a multiphase liquid condensate. Nat. Rev. Mol. Cell Biol., 22, 165–182.

6. Gorenstein, C. and Warner, J.R. (1977) Synthesis and turnover of ribosomal proteins in the absence of 60S subunit assembly in Saccharomyces cerevisiae. Mol. Gen. Genet., 157, 327–332.

7. Warner, J.R. (1977) In the absence of ribosomal RNA synthesis, the ribosomal proteins of HeLa cells are synthesized normally and degraded rapidly. J. Mol. Biol., 115, 315–333.

8. Sung, M.K., Porras-Yakushi, T.R., Reitsma, J.M., Huber, F.M., Sweredoski, M.J., Hoelz, A., Hess, S. and Deshaies, R.J. (2016) A conserved quality-control pathway that mediates degradation of unassembled ribosomal proteins. Elife, 5.

9. Sung, M.K., Reitsma, J.M., Sweredoski, M.J., Hess, S. and Deshaies, R.J. (2016) Ribosomal proteins produced in excess are degraded by the ubiquitin-proteasome system. Mol. Biol. Cell, 27, 2642–2652.

10. Juli, G., Gismondi, A., Monteleone, V., Caldarola, S., Iadevaia, V., Aspesi, A., Dianzani, I., Proud, C.G. and Loreni, F. (2016) Depletion of ribosomal protein S19 causes a reduction of rRNA synthesis. Sci Rep, 6, 35026.

11. O’Donohue, M.F., Choesmel, V., Faubladier, M., Fichant, G. and Gleizes, P.E. (2010) Functional dichotomy of ribosomal proteins during the synthesis of mammalian 40S ribosomal subunits. J. Cell Biol., 190, 853–866.

12. Venturi, G. and Montanaro, L. (2020) How Altered Ribosome Production Can Cause or Contribute to Human Disease: The Spectrum of Ribosomopathies. Cells, 9.

13. Da Costa, L., Leblanc, T. and Mohandas, N. (2020) Diamond-Blackfan anemia. Blood, 136, 1262–1273.

14. Fonseca, B.D., Zakaria, C., Jia, J.J., Graber, T.E., Svitkin, Y., Tahmasebi, S., Healy, D., Hoang, H.D., Jensen, J.M., Diao, I.T. et al. (2015) La-related Protein 1 (LARP1) Represses Terminal Oligopyrimidine (TOP) mRNA Translation Downstream of mTOR Complex 1 (mTORC1). J. Biol. Chem., 290, 15996–16020.

15. Jia, J.J., Lahr, R.M., Solgaard, M.T., Moraes, B.J., Pointet, R., Yang, A.D., Celucci, G., Graber, T.E., Hoang, H.D., Niklaus, M.R. et al. (2021) mTORC1 promotes TOP mRNA translation through site-specific phosphorylation of LARP1. Nucleic Acids Res., 49, 3461–3489.

16. Mayer, C., Zhao, J., Yuan, X. and Grummt, I. (2004) mTOR-dependent activation of the transcription factor TIF-IA links rRNA synthesis to nutrient availability. Genes Dev., 18, 423–434.

17. Pakos-Zebrucka, K., Koryga, I., Mnich, K., Ljujic, M., Samali, A. and Gorman, A.M. (2016) The integrated stress response. EMBO Rep., 17, 1374–1395.

18. Ranu, R.S. and London, I.M. (1976) Regulation of protein synthesis in rabbit reticulocyte lysates: purification and initial characterization of the cyclic 3’:5’-AMP independent protein kinase of the heme-regulated translational inhibitor. Proc. Natl. Acad. Sci. U. S. A., 73, 4349–4353.

19. Levin, D.H., Petryshyn, R. and London, I.M. (1980) Characterization of double-stranded-RNA-activated kinase that phosphorylates alpha subunit of eukaryotic initiation factor 2 (eIF-2 alpha) in reticulocyte lysates. Proc. Natl. Acad. Sci. U. S. A., 77, 832–836.

20. Shi, Y., Vattem, K.M., Sood, R., An, J., Liang, J., Stramm, L. and Wek, R.C. (1998) Identification and characterization of pancreatic eukaryotic initiation factor 2 alpha-subunit kinase, PEK, involved in translational control. Mol. Cell. Biol., 18, 7499–7509.

21. Sood, R., Porter, A.C., Olsen, D.A., Cavener, D.R. and Wek, R.C. (2000) A mammalian homologue of GCN2 protein kinase important for translational control by phosphorylation of eukaryotic initiation factor-2alpha. Genetics, 154, 787–801.

22. Aulas, A., Fay, M.M., Lyons, S.M., Achorn, C.A., Kedersha, N., Anderson, P. and Ivanov, P. (2017) Stress-specific differences in assembly and composition of stress granules and related foci. J. Cell Sci., 130, 927–937.

23. Patel, K.S., Pandey, P.K., Martin-Ramos, P., Corns, W.T., Varol, S., Bhattacharya, P. and Zhu, Y. (2023) A review on arsenic in the environment: bio-accumulation, remediation, and disposal. RSC Adv, 13, 14914–14929.

24. Henras, A.K., Plisson-Chastang, C., O’Donohue, M.F., Chakraborty, A. and Gleizes, P.E. (2015) An overview of pre-ribosomal RNA processing in eukaryotes. Wiley interdisciplinary reviews. RNA, 6, 225–242.

25. Gurney, T., Jr. (1985) Characterization of mouse 45S ribosomal RNA subspecies suggests that the first processing cleavage occurs 600 +/- 100 nucleotides from the 5’ end and the second 500 +/- 100 nucleotides from the 3’ end of a 13.9 kb precursor. Nucleic Acids Res., 13, 4905–4919.

26. Tiollais, P., Galibert, F. and Boiron, M. (1971) Evidence for the existence of several molecular species in the "45S fraction" of mammalian ribosomal precursor RNA. Proc. Natl. Acad. Sci. U. S. A., 68, 1117–1120.

27. Mullineux, S.T. and Lafontaine, D.L. (2012) Mapping the cleavage sites on mammalian pre-rRNAs: where do we stand? Biochimie, 94, 1521–1532.

28. Lesniczak-Staszak, M., Pietras, P., Rucinski, M., Johnston, R., Sowinski, M., Andrzejewska, M., Nowicki, M., Gowin, E., Lyons, S.M., Ivanov, P. et al. (2024) Stress granule-mediated sequestration of EGR1 mRNAs correlates with lomustine-induced cell death prevention. J. Cell Sci., 137.

29. James, M.J. and Zomerdijk, J.C. (2004) Phosphatidylinositol 3-kinase and mTOR signaling pathways regulate RNA polymerase I transcription in response to IGF-1 and nutrients. J. Biol. Chem., 279, 8911–8918.

30. Brangwynne, C.P., Eckmann, C.R., Courson, D.S., Rybarska, A., Hoege, C., Gharakhani, J., Julicher, F. and Hyman, A.A. (2009) Germline P granules are liquid droplets that localize by controlled dissolution/condensation. Science, 324, 1729–1732.

31. Zeng, M., Xiao, F., Zhong, X., Jin, F., Guan, L., Wang, A., Liu, X. and Zhong, C. (2013) Reactive oxygen species play a central role in hexavalent chromium-induced apoptosis in Hep3B cells without the functional roles of p53 and caspase-3. Cell. Physiol. Biochem., 32, 279–290.

32. Liu, K.J. and Shi, X. (2001) In vivo reduction of chromium (VI) and its related free radical generation. Mol. Cell. Biochem., 222, 41–47.

33. Meaza, I., Williams, A.R., Wise, S.S., Lu, H. and Wise, J.P., Sr. (2024) Carcinogenic Mechanisms of Hexavalent Chromium: From DNA Breaks to Chromosome Instability and Neoplastic Transformation. Curr Environ Health Rep, 11, 484–546.

34. Wang, Z. and Yang, C. (2023) Epigenetic and epitranscriptomic mechanisms of chromium carcinogenesis. Adv. Pharmacol., 96, 241–265.

35. Stannard, L.M., Doherty, A., Chapman, K.E., Doak, S.H. and Jenkins, G.J. (2024) Multi-endpoint analysis of cadmium chloride-induced genotoxicity shows role for reactive oxygen species and p53 activation in DNA damage induction, cell cycle irregularities, and cell size aberrations. Mutagenesis, 39, 13–23.

36. Skipper, A., Sims, J.N., Yedjou, C.G. and Tchounwou, P.B. (2016) Cadmium Chloride Induces DNA Damage and Apoptosis of Human Liver Carcinoma Cells via Oxidative Stress. Int J Environ Res Public Health, 13.

37. Wallace, D.R., Spandidos, D.A., Tsatsakis, A., Schweitzer, A., Djordjevic, V. and Djordjevic, A.B. (2019) Potential interaction of cadmium chloride with pancreatic mitochondria: Implications for pancreatic cancer. Int. J. Mol. Med., 44, 145–156.

38. Ursu, G.M., Krawic, C. and Zhitkovich, A. (2025) Nuclear SUMOylation and Proteotoxic Stress Responses to Metals with Different Ligand Preferences. Chem. Res. Toxicol., 38, 942–953.

39. Gingras, A.C., Gygi, S.P., Raught, B., Polakiewicz, R.D., Abraham, R.T., Hoekstra, M.F., Aebersold, R. and Sonenberg, N. (1999) Regulation of 4E-BP1 phosphorylation: a novel two-step mechanism. Genes Dev., 13, 1422–1437.

40. Pause, A., Belsham, G.J., Gingras, A.C., Donze, O., Lin, T.A., Lawrence, J.C., Jr. and Sonenberg, N. (1994) Insulin-dependent stimulation of protein synthesis by phosphorylation of a regulator of 5’-cap function. Nature, 371, 762–767.

41. Yao, R.W., Xu, G., Wang, Y., Shan, L., Luan, P.F., Wang, Y., Wu, M., Yang, L.Z., Xing, Y.H., Yang, L. et al. (2019) Nascent Pre-rRNA Sorting via Phase Separation Drives the Assembly of Dense Fibrillar Components in the Human Nucleolus. Mol. Cell, 76, 767–783 e711.

42. Kuhn, A. and Grummt, I. (1989) 3’-end formation of mouse pre-rRNA involves both transcription termination and a specific processing reaction. Genes Dev., 3, 224–231.

43. Grummt, I., Rosenbauer, H., Niedermeyer, I., Maier, U. and Ohrlein, A. (1986) A repeated 18 bp sequence motif in the mouse rDNA spacer mediates binding of a nuclear factor and transcription termination. Cell, 45, 837–846.

44. Buratowski, S. (2005) Connections between mRNA 3’ end processing and transcription termination. Curr. Opin. Cell Biol., 17, 257–261.

45. West, S., Gromak, N. and Proudfoot, N.J. (2004) Human 5’ --> 3’ exonuclease Xrn2 promotes transcription termination at co-transcriptional cleavage sites. Nature, 432, 522–525.

46. Lopez Martinez, D. and Svejstrup, J.Q. (2025) Mechanisms of RNA Polymerase II Termination at the 3’-End of Genes. J. Mol. Biol., 437, 168735.

47. Birse, C.E., Minvielle-Sebastia, L., Lee, B.A., Keller, W. and Proudfoot, N.J. (1998) Coupling termination of transcription to messenger RNA maturation in yeast. Science, 280, 298–301.

48. Connelly, S. and Manley, J.L. (1988) A functional mRNA polyadenylation signal is required for transcription termination by RNA polymerase II. Genes Dev., 2, 440–452.

49. El Hage, A., Koper, M., Kufel, J. and Tollervey, D. (2008) Efficient termination of transcription by RNA polymerase I requires the 5’ exonuclease Rat1 in yeast. Genes Dev., 22, 1069–1081.

50. Petfalski, E., Winz, M.L., Grelewska-Nowotko, K., Turowski, T.W. and Tollervey, D. (2025) Multiple mechanisms of termination modulate the dynamics of RNAPI transcription. Cell Rep, 44, 115325.

51. Kass, S., Tyc, K., Steitz, J.A. and Sollner-Webb, B. (1990) The U3 small nucleolar ribonucleoprotein functions in the first step of preribosomal RNA processing. Cell, 60, 897–908.

52. Kufel, J., Dichtl, B. and Tollervey, D. (1999) Yeast Rnt1p is required for cleavage of the pre-ribosomal RNA in the 3’ ETS but not the 5’ ETS. RNA, 5, 909–917.

53. Kempers-Veenstra, A.E., Oliemans, J., Offenberg, H., Dekker, A.F., Piper, P.W., Planta, R.J. and Klootwijk, J. (1986) 3’-End formation of transcripts from the yeast rRNA operon. EMBO J, 5, 2703–2710.

54. Silva, S., Homolka, D. and Pillai, R.S. (2017) Characterization of the mammalian RNA exonuclease 5/NEF-sp as a testis-specific nuclear 3’ --> 5’ exoribonuclease. RNA, 23, 1385–1392.

55. Peculis, B.A. (1997) The sequence of the 5’ end of the U8 small nucleolar RNA is critical for 5.8S and 28S rRNA maturation. Mol. Cell. Biol., 17, 3702–3713.

56. Langhendries, J.L., Nicolas, E., Doumont, G., Goldman, S. and Lafontaine, D.L. (2016) The human box C/D snoRNAs U3 and U8 are required for pre-rRNA processing and tumorigenesis. Oncotarget, 7, 59519–59534.

57. Levy, S., Avni, D., Hariharan, N., Perry, R.P. and Meyuhas, O. (1991) Oligopyrimidine tract at the 5’ end of mammalian ribosomal protein mRNAs is required for their translational control. Proc. Natl. Acad. Sci. U. S. A., 88, 3319–3323.

58. Avni, D., Biberman, Y. and Meyuhas, O. (1997) The 5’ terminal oligopyrimidine tract confers translational control on TOP mRNAs in a cell type- and sequence context-dependent manner. Nucleic Acids Res., 25, 995–1001.

59. Patursky-Polischuk, I., Stolovich-Rain, M., Hausner-Hanochi, M., Kasir, J., Cybulski, N., Avruch, J., Ruegg, M.A., Hall, M.N. and Meyuhas, O. (2009) The TSC-mTOR pathway mediates translational activation of TOP mRNAs by insulin largely in a raptor- or rictor-independent manner. Mol. Cell. Biol., 29, 640–649.

60. Sprenkle, N.T., Sims, S.G., Sanchez, C.L. and Meares, G.P. (2017) Endoplasmic reticulum stress and inflammation in the central nervous system. Mol Neurodegener, 12, 42.

61. Bell, M.C., Meier, S.E., Ingram, A.L. and Abisambra, J.F. (2016) PERK-opathies: An Endoplasmic Reticulum Stress Mechanism Underlying Neurodegeneration. Curr. Alzheimer Res., 13, 150–163.

62. Oakes, S.A. and Papa, F.R. (2015) The role of endoplasmic reticulum stress in human pathology. Annu Rev Pathol, 10, 173–194.

63. Halliday, M., Hughes, D. and Mallucci, G.R. (2017) Fine-tuning PERK signaling for neuroprotection. J. Neurochem., 142, 812–826.

64. Moreno, J.A., Halliday, M., Molloy, C., Radford, H., Verity, N., Axten, J.M., Ortori, C.A., Willis, A.E., Fischer, P.M., Barrett, D.A. et al. (2013) Oral treatment targeting the unfolded protein response prevents neurodegeneration and clinical disease in prion-infected mice. Sci Transl Med, 5, 206ra138.

65. Radford, H., Moreno, J.A., Verity, N., Halliday, M. and Mallucci, G.R. (2015) PERK inhibition prevents tau-mediated neurodegeneration in a mouse model of frontotemporal dementia. Acta Neuropathol., 130, 633–642.

66. Celardo, I., Costa, A.C., Lehmann, S., Jones, C., Wood, N., Mencacci, N.E., Mallucci, G.R., Loh, S.H. and Martins, L.M. (2016) Mitofusin-mediated ER stress triggers neurodegeneration in pink1/parkin models of Parkinson’s disease. Cell Death Dis, 7, e2271.

67. Halliday, M., Radford, H., Sekine, Y., Moreno, J., Verity, N., le Quesne, J., Ortori, C.A., Barrett, D.A., Fromont, C., Fischer, P.M., et al. (2015) Partial restoration of protein synthesis rates by the small molecule ISRIB prevents neurodegeneration without pancreatic toxicity. Cell Death Dis, 6, e1672.

68. Salnikow, K. and Zhitkovich, A. (2008) Genetic and epigenetic mechanisms in metal carcinogenesis and cocarcinogenesis: nickel, arsenic, and chromium. Chem. Res. Toxicol., 21, 28–44.

69. Reynolds, R.C., Montgomery, P.O. and Hughes, B. (1964) Nucleolar "Caps" Produced by Actinomycin D. Cancer Res., 24, 1269–1277.

70. Tchelidze, P., Benassarou, A., Kaplan, H., O’Donohue, M.F., Lucas, L., Terryn, C., Rusishvili, L., Mosidze, G., Lalun, N. and Ploton, D. (2017) Nucleolar sub-compartments in motion during rRNA synthesis inhibition: Contraction of nucleolar condensed chromatin and gathering of fibrillar centers are concomitant. PLoS One, 12, e0187977.

