## Supplemental Figures for "Metal stress uncouples early pre-rRNA processing from ISR activation and reveals flexible checkpoints in human ribosome biogenesis"

**Supplemental Information**

**Supplementary Figure 1.** Quantification of HeLa, HAP1, and U2OS Northern blots (n=3). (**A**) 5'ETS quantification for control and $AsO_2$ treated cells grouped by cell line. Data analyzed by multiple paired t-tests (* $P < 0.05$). All cell lines indicate similar trends of precursor accumulation (**B**) ITS1 quantification for the three cell lines indicate more variability within these precursors, notably U2OS cells exhibiting substantial decrease in key species. (**C**) ITS2 quantification shows similar trending of all cell lines. (**D**) 3'ETS quantification shows clearance of major precursors across cell lines.

**Supplementary Figure 2.** Total quantification of all precursors for Tun, Tg, and DTT treated cells from ITS1 probe (n=3). Data analyzed by one way ANOVA with Dunnett's multiple comparisons post-hoc test. (**A**) 47S/47S-C/45S/46S quantification of the ER stressors show some modulation of abundance across concentrations tested. (*ns* $P > 0.05$, * $P < 0.05$) (**B**) 43S precursor quantification significantly down at the lowest concentration of Tun, though unperturbed in other conditions. (**C**) 26S quantification variably decreases abundance upon treatment of all stressors, though notably follows a dose dependent decreased trend for DTT. (* $P < 0.05$, ** $P < 0.01$, *** $P < 0.001$) (**D**) 21S precursor quantification reveals no significant changes in accumulation. (**E**) 18S-E quantification for ITS1 probe shows no significant changes upon treatment of ER stressors.

**Supplementary Figure 3.** Total quantification of remaining precursors captured with ITS2 probe and precursors captured with 3'ETS probe treated with ER Stressors (n=3). Data analyzed by one way ANOVA with Dunnett's multiple comparisons post-hoc test (*ns* $P > 0.05$). (**A**) 47S/47S-C/45S/46S ITS2 quantification indicates no significant change in levels of the largest precursors. (**B**) ITS2 12S levels are unperturbed across treatments and concentrations compared to control.

(**C**) 3'ETS 47S/46S species levels are not significantly changed across test samples. (**D**) 28S-L 3'ETS quantification reveals a significant increase in species level at the highest concentration of Tun, though no significance elsewhere in the samples tested (* $P < 0.05$).

**Supplementary Figure 4.** Total quantification of all precursors for heavy metal treated cells from ITS1 probe (n=3). Data analyzed by one way ANOVA with Dunnett's multiple comparisons post-hoc test (*ns* $P > 0.05$). (**A**) ITS1 47S/47S-C/45S/46S quantification shows $CrO_3$ significantly depletes the largest precursors at all concentrations tested (* $P < 0.05$). (**B**) 43S ITS1 precursor is depleted significantly upon chromium treatment at all concentrations (** $P < 0.01$ ). (**C**) 26S precursor detected by ITS1 probe is significantly down upon all treatments and concentrations tested save the lowest concentration (100 µM) of arsenite (*** $P < 0.001$). (**D**) 21S/21S-C species from ITS1 is significantly down upon all chromium treatments (**** $P < 0.0001$), and dose dependently decrease upon $CdCl_2$ treatment. (**E**) 18S-E species detected by ITS1 probe dose dependently decrease in response to $AsO_2$ and are depleted across all concentrations of $CrO_3$ tested.

**Supplementary Figure 5.** Heavy metal total quantification of remaining precursors captured with ITS2 and 3'ETS probe (n=3). Data analyzed by one way ANOVA with Dunnett's multiple comparisons post-hoc test (*ns* $P > 0.05$). (**A**) ITS2 47S/47S-C/45S/46S quantification shows no significant change in accumulation of the largest rRNA precursor species in response to heavy metal treatment. (**B**) 12S ITS2 levels are significantly decreased by the highest concentration of $CdCl_2$ (* $P < 0.05$), and qualitatively decrease dose dependently. (**C**) 3'ETS 47S/46S precursor levels are significantly decreased by all concentrations of $CrO_3$ tested (**** $P < 0.0001$). (**D**) 3'ETS 28S-L levels are unperturbed by the heavy metals at tested concentrations.

**Supplementary Figure 6.** Northern blot and quantification of major precursors from heavy metal titrations in HeLa cells (n=3). (**A**) RNA from HeLa cells treated with titrations of $AsO_2$ (100, 300, 500 µM), $CrO_3$ (100, 300, 500 µM), and CdCl2 (100, 300, 500 µM) run on a Northern blot. Ethidium bromide (EtBr) images of mature 28S and 18S rRNA included for loading. (**B**) Chromium significantly depletes the largest 5'ETS captured precursors (47S/47S-C) at the highest concentration tested (*ns* $P > 0.05$, * $P < 0.05$). (**C**) Heavy metals have no significant effect on 34S species accumulation, though AsO2 and CdCl2 both exhibit qualitative dose dependent increases in accumulation. (**D**) No significant changes in accumulation of 30S (34S*) species probed with ITS1 are apparent. (**E**) 32S accumulation is unchanged in the presence of heavy metal treatment across all compounds and concentrations.

### Supplemental Figure 1

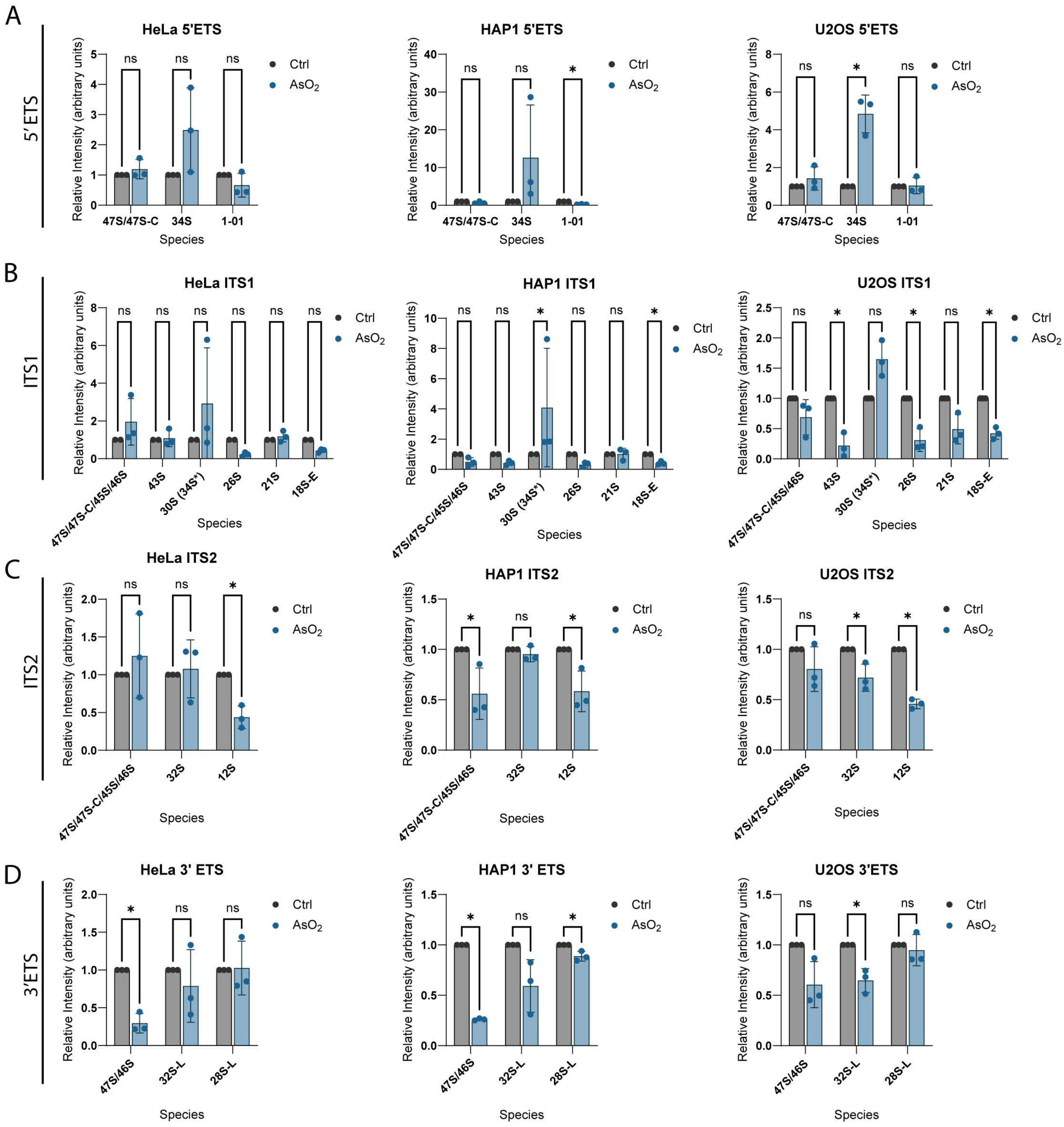

Supplementary Figure 2

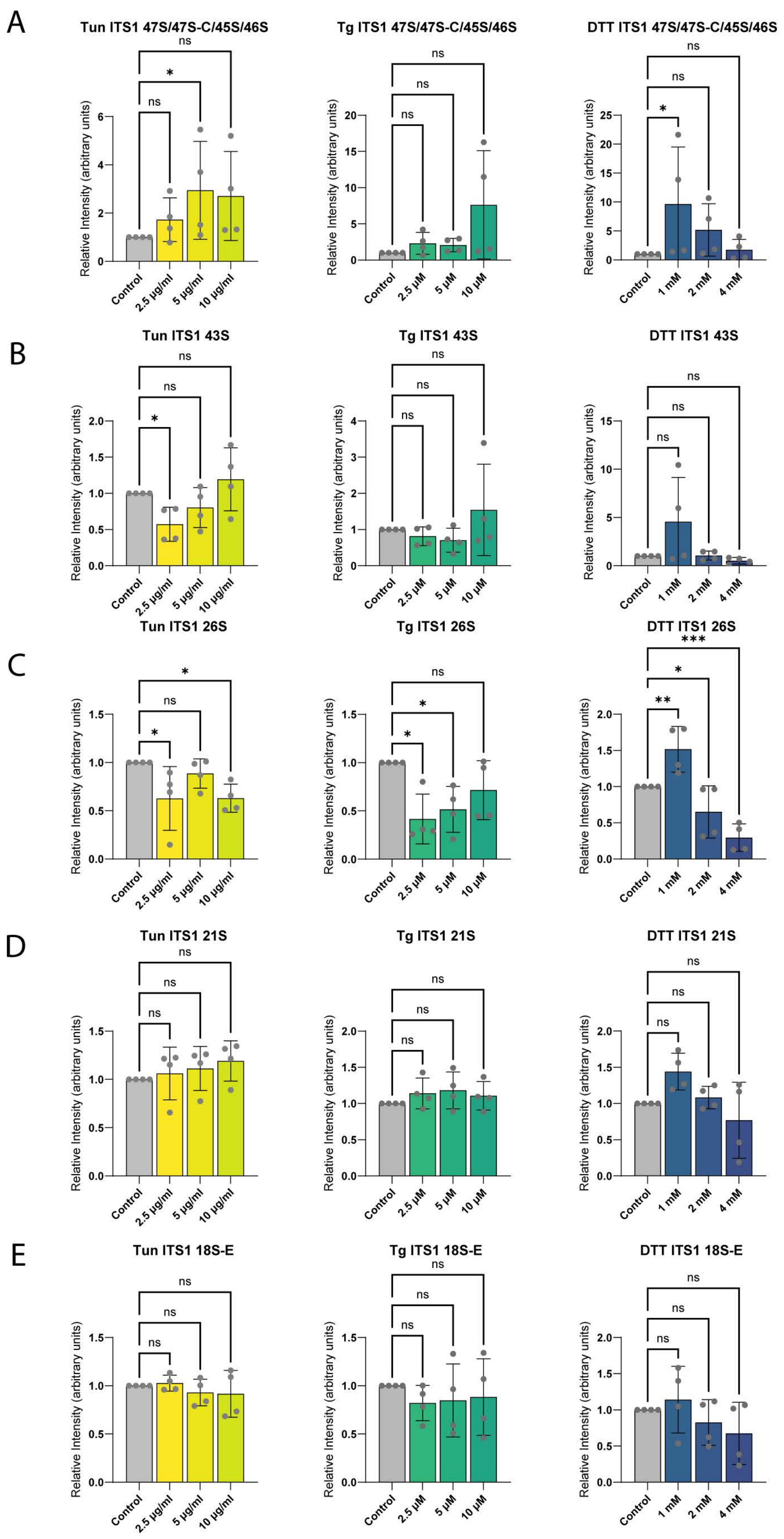

### Supplementary Figure 3

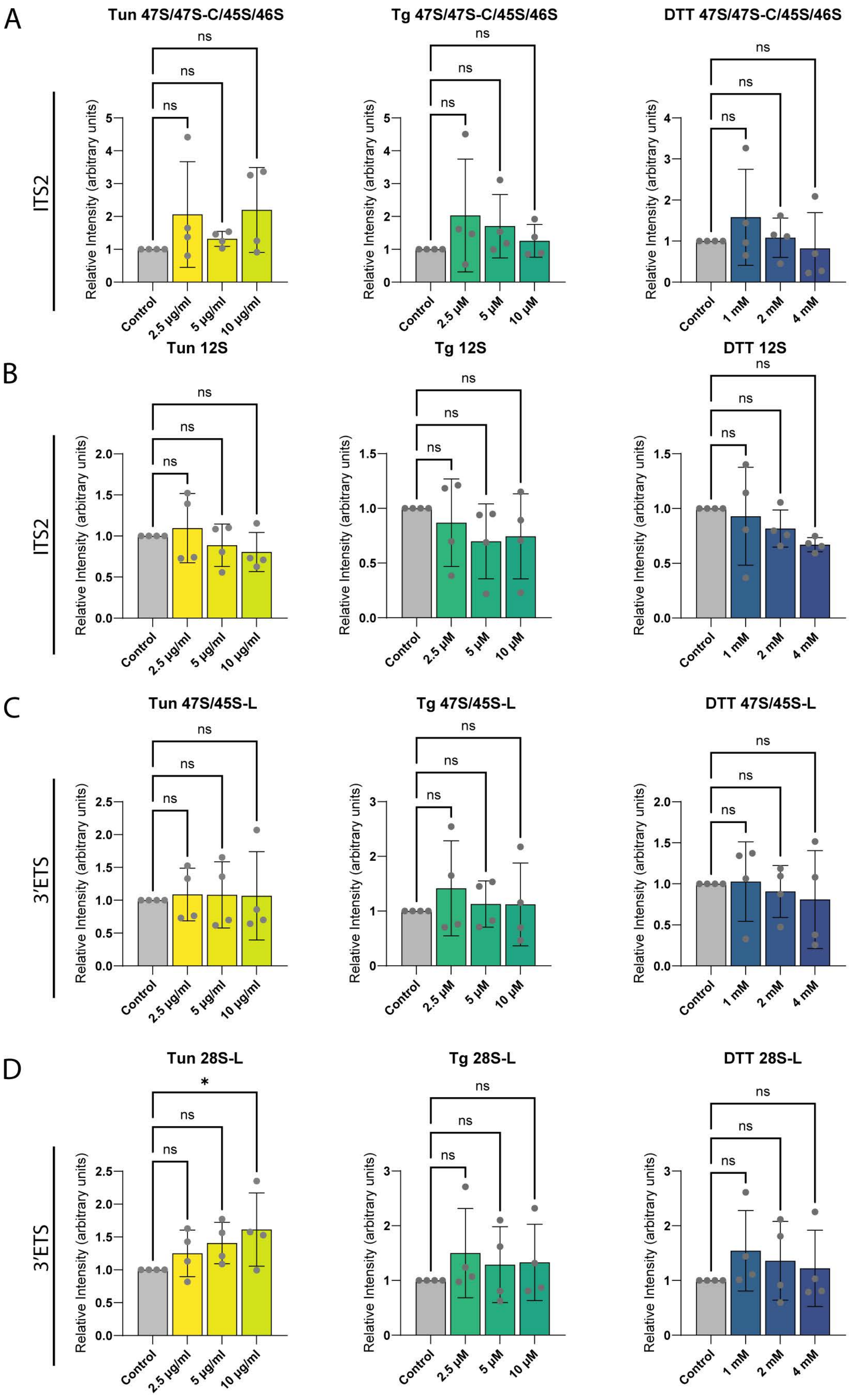

Supplementary Figure 4

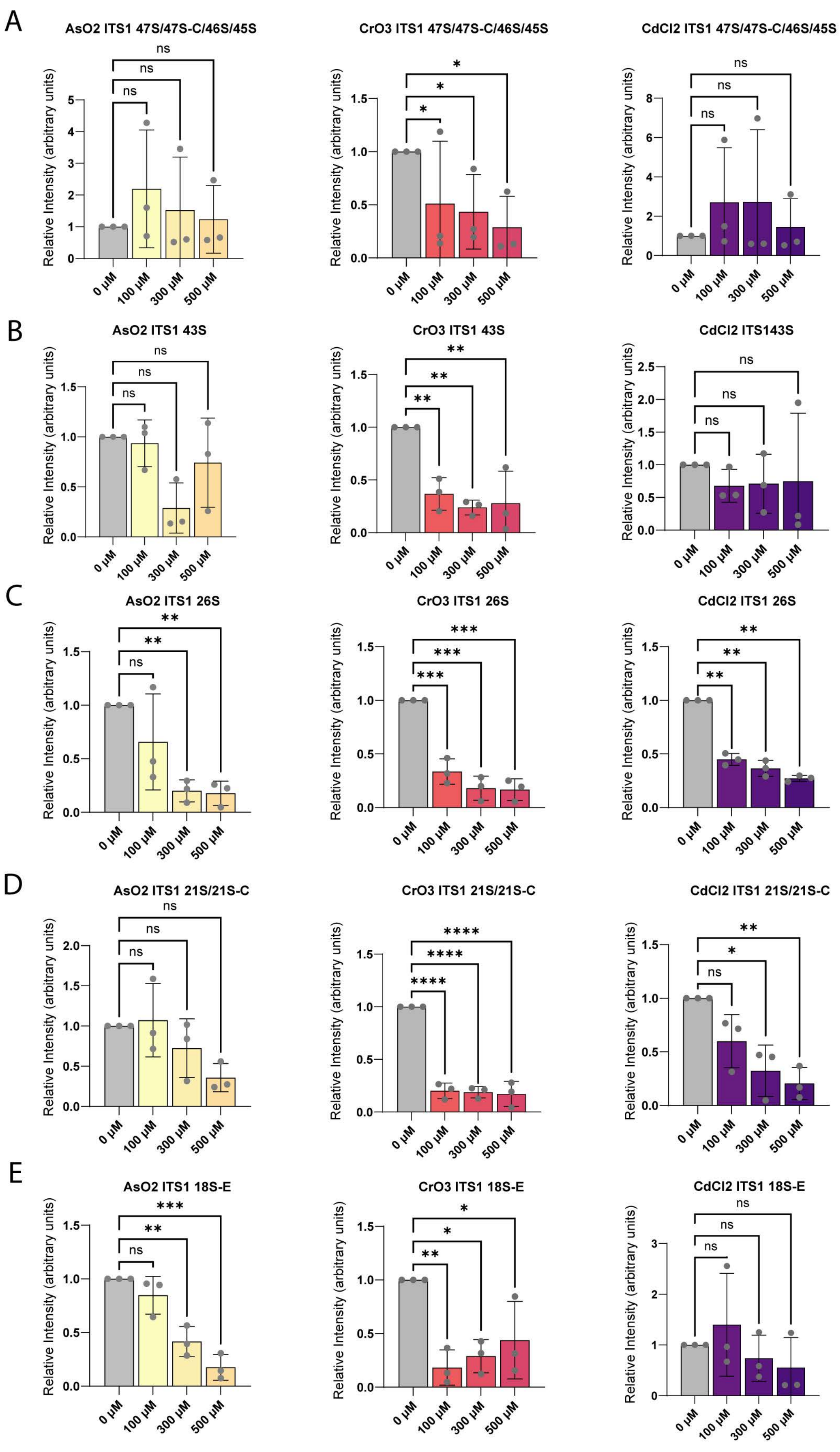

Supplementary Figure 5

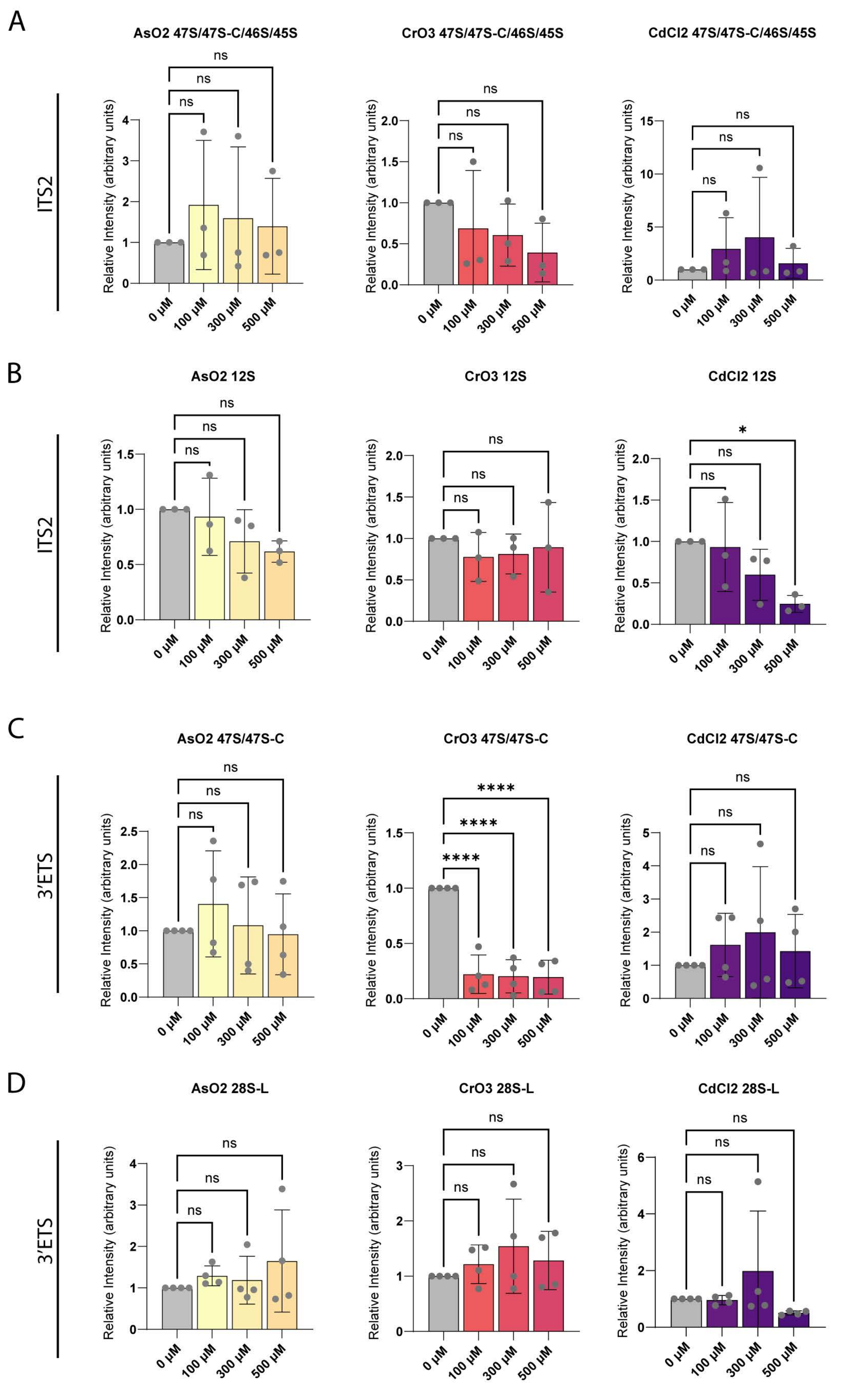

### Supplemental Figure 6

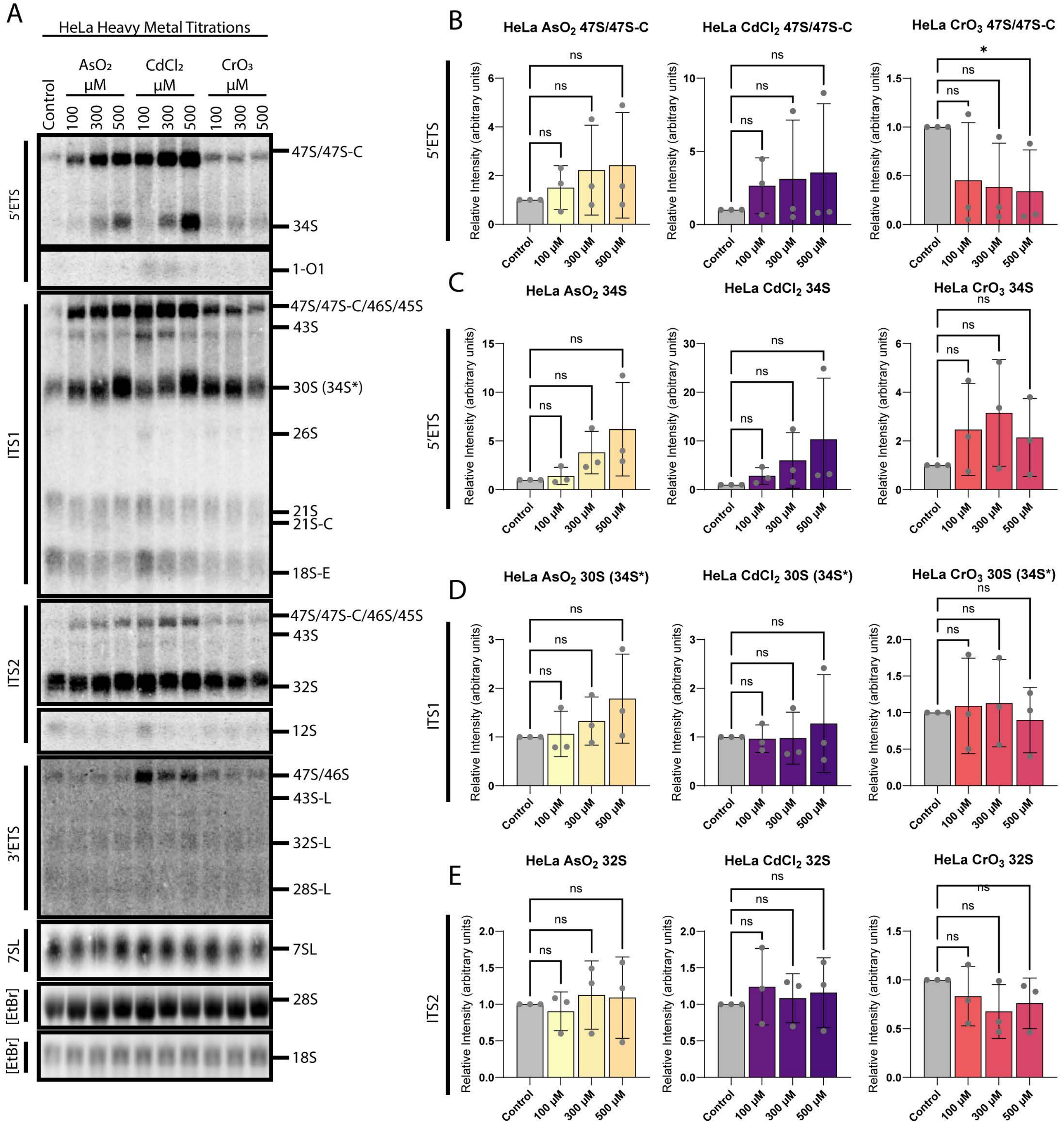
